# Neuroprotective derivatives of tacrine that target NMDA receptor and acetyl cholinesterase - Design, synthesis and biological evaluation

**DOI:** 10.1101/2021.01.18.427058

**Authors:** Chandran Remya, K.V. Dileep, Eeda Koti Reddy, Kumar Mantosh, Kesavan Lakshmi, Reena Sarah Jacob, Ayyiliyath M Sajith, E.Jayadevi Variyar, Shaik Anwar, Kam Y. J. Zhang, C. Sadasivan, R. V. Omkumar

## Abstract

The complex and multifactorial nature of neuropsychiatric diseases demands multi-target drugs that can intervene with various sub-pathologies underlying disease progression. Targeting the impairments in cholinergic and glutamatergic neurotransmissions with small molecules has been suggested as one of the potential disease-modifying approaches for Alzheimer’s disease (AD). Tacrine, apotent inhibitor of acetylcholinesterase (AChE) is the first FDA approved drug for the treatment of AD. Tacrine is also a low affinity antagonist of N-methyl-D-aspartate receptor (NMDAR). However, tacrine was withdrawn from its clinical use later due to its hepatotoxicity. With an aim to develop novel high affinity multi-target directed ligands (MTDLs) against AChE and NMDAR, with reduced hepatotoxicity, we performed *in silico* structure-based modifications on tacrine, chemical synthesis of the derivatives and *in vitro* validation of their activities. Nineteen such derivatives showed inhibition with IC_50_ values in the range of 18.53±2.09 to 184.09±19.23 nM against AChE and 0.27±0.05 to 38.84±9.64 μM against NMDAR. Some of the selected compounds also protected rat primary cortical neurons from glutamate induced excitotoxicity. Two of the tacrine derived MTDLs, 201 and 208 exhibited *in vivo* efficacy in rats by protecting against behavioral impairment induced by administration of the excitotoxic agent, monosodium glutamate. Additionally, several of these synthesized compounds also exhibited promising inhibitory activities against butyrylcholinesterase and β-secretase. Given the therapeutic potential of MTDLs in disease-modifying therapy, our studies revealed several promising MTDLs of which 201 appears to be a potential candidate for immediate preclinical and clinical evaluations.

## Introduction

Alterations in the levels of various neurotransmitters and functioning of neuronal networks in the brain lead to neuropsychiatric disorders. Alzheimer’s disease (AD) is a condition in which various factors such as impairment in cholinergic and glutamatergic signaling, toxicity due to accumulation of Aβ peptides and hyperphosphorylated tau proteins, neuroinflammation, metal dyshomeostasis, oxidative stress, mitochondrial dysfunction and genetic predisposition contribute to the pathological events leading to cognitive decline and neurodegeneration [1–3]. Correlation between cholinergic dysfunction and AD progression prompted the identification of several potential disease-modifying agents such as acetylcholinesterase inhibitors (AChEIs) and acetylcholine receptor agonists for improving cholinergic functions [4–10]. AChEIs such as donepezil [11], galanthamine [12] and rivastigmine [13] are currently being used for the treatment of moderate to severe AD. Though these drugs are beneficial in improving cognitive and behavioral symptoms, they do not prevent the process of neurodegeneration completely.

Neuronal loss that occurs in the brain is an underlying factor for AD [14, 15] and for other neurodegenerative diseases [16]. Abnormal release of glutamate and/or deficiency of glutamate uptake mechanisms result in the accumulation of extracellular glutamate leading to neuronal apoptosis, a process termed as excitotoxicity, that happens by the overactivation of the *N*-methyl-*D*-aspartate type glutamate receptor (NMDAR) [17–19]. NMDAR antagonists are suggested to be therapeutic agents for this condition [19, 20]. Excitotoxicity triggered by Aβ peptides and tau proteins have been prevented by NMDAR antagonists like memantine and ifenprodil [21–23]. Memantine is one of the FDA approved drugs used for the treatment of moderate to severe AD [24]. According to the current clinical data, combination therapy with memantine and AChEIs produces benefits in all stages of AD than the monotherapies [25].

In conditions like oxygen-glucose deprivation in brain slices [26] and in NMDA-induced excitotoxicity in cultured neurons [27], the role of NMDAR as a potential target for neuroprotection has already been demonstrated. Variations in NMDAR activity are associated with ischemic conditions [28], stroke [29], traumatic brain injury (TBI) [30], glioma [31] and neuropsychiatric diseases [32, 33], suggesting the therapeutic potential of NMDAR modulators in these conditions. The involvement of NMDARs in pain circuitries makes it a potential target for analgesic drugs [34]. However, many NMDAR antagonists currently in clinical use either have insufficient efficacy or have undesirable side effects. Hence, new and better antagonists are necessary for treating neurological diseases.

Tacrine, a potent inhibitor of AChE [35] is the first FDA approved drug for the treatment of AD but had hepatotoxicity that led to its withdrawal from its clinical use [36]. Interestingly, tacrine was also reported as a weak antagonist of NMDAR [37]. Hence, tacrine is unlikely to cause NMDAR inhibition at its therapeutic dose required to achieve AChE inhibition. This also makes tacrine an unsuitable candidate for the treatment of other neurological conditions such as stroke and traumatic brain injury where NMDAR antagonists would be useful. Chemical modification of tacrine may help to improve its inhibitory potency towards NMDAR. This may also permit reducing the dosage so that hepatotoxicity could be brought within safety limits. Despite its hepatotoxicity, tacrine structure has been successfully used in medicinal chemistry for designing hybrids and multi-target directed ligands (MTDLs) [38–48].

Due to the complex etiology and multi-faceted nature of AD, use of MTDLs has been suggested as a promising treatment strategy. Compared to the mono and combination therapies, MTDLs have advantages [49] such as lower probabilities of drug-drug interactions, reduced off-target interactions, wider therapeutic window and improved safety profiles [50, 51]. Though several MTDLs were shown to be effective *in vitro*, success rate in the preclinical/clinical stages have been highly limited. By combining molecules such as donepezil, galantamine, rivastigmine and tacrine with each other and with other chemical entities, several MTDLs have been suggested [52].

In the current study, with an aim to propose novel, high affinity and less hepatotoxic tacrine derived MTDLs that can modulate the dysfunctions in cholinergic and glutamatergic systems, we systematically designed, synthesized and evaluated a series of MTDLs against the two promising drug targets, AChE and NMDAR. Effects of MTDLs on excitotoxic conditions in rat primary cortical neurons were also studied. Neuroprotective effects of selected MTDLs were also assessed using behavioral study of rats with the help of Morris water maze test (MWM).

## Results

### Molecular modeling studies

#### Coarse-grained modeling provided h-NMDARs with multiple receptor conformations

High-resolution crystal structures of quasi-independent domains of NMDAR with different allosteric inhibitors have been reported [53, 54]. However, structures representing the open and closed conformations of the channels were reported at moderate resolutions. Missing residues and side chains in these structures made them inappropriate for the molecular modeling studies. Reports from literature suggested that tacrine binds in the channel of NMDAR [55–60]. To decrypt the mode of binding of tacrine in the channel pore of NMDAR, we assumed that the structural dynamics of NMDAR have to be addressed. The GluN1/GluN2B subtype of NMDAR of *Xenopus laevis* possessed high sequence similarity (≥ 90%) to h-GluN1/GluN2B (h-NMDAR). Hence, h-NMDAR was modelled by choosing crystal structure of GluN1/GluN2B heteromer of *Xenopus laevis* in complex with MK-801 as the template. The missing regions in the template structure, such as loops were modelled and energy minimized. The lowest energy conformation was taken as the model for further studies. As expected the modeled structure showed high similarity to the template structure.

Since all atom MD simulation was computationally very expensive, coarse-grained modeling technique was used to probe the dynamics of h-NMDAR. The observed structural dynamics in the coarse-grained modeling were assumed to be representing various biological functional states such as open and closed conformations of the channel. The NMA and ENM simulations retrieved 339 output structures, which were gathered and checked for the global root mean square deviation (RMSD) by superimposing them with the modeled h-NMDAR. The superimposition studies revealed that RMSD varies from ~ 0.1 to 4 Å. Based on RMSD, the structures were classified into four clusters. The cluster-1 contains structures with RMSD 0.1 to 1Å, cluster-2 contains structures with RMSD 1-2 Å, cluster-3 with RMSD 2-3Å and cluster-4 with RMSD 3-4Å (Fig. 1A). Since the template structure is in the open conformation, all the structures in cluster-1 were assumed to be representing an open conformation. Candidate structures were selected from each cluster and were used for the docking studies.

**Figure 1:**
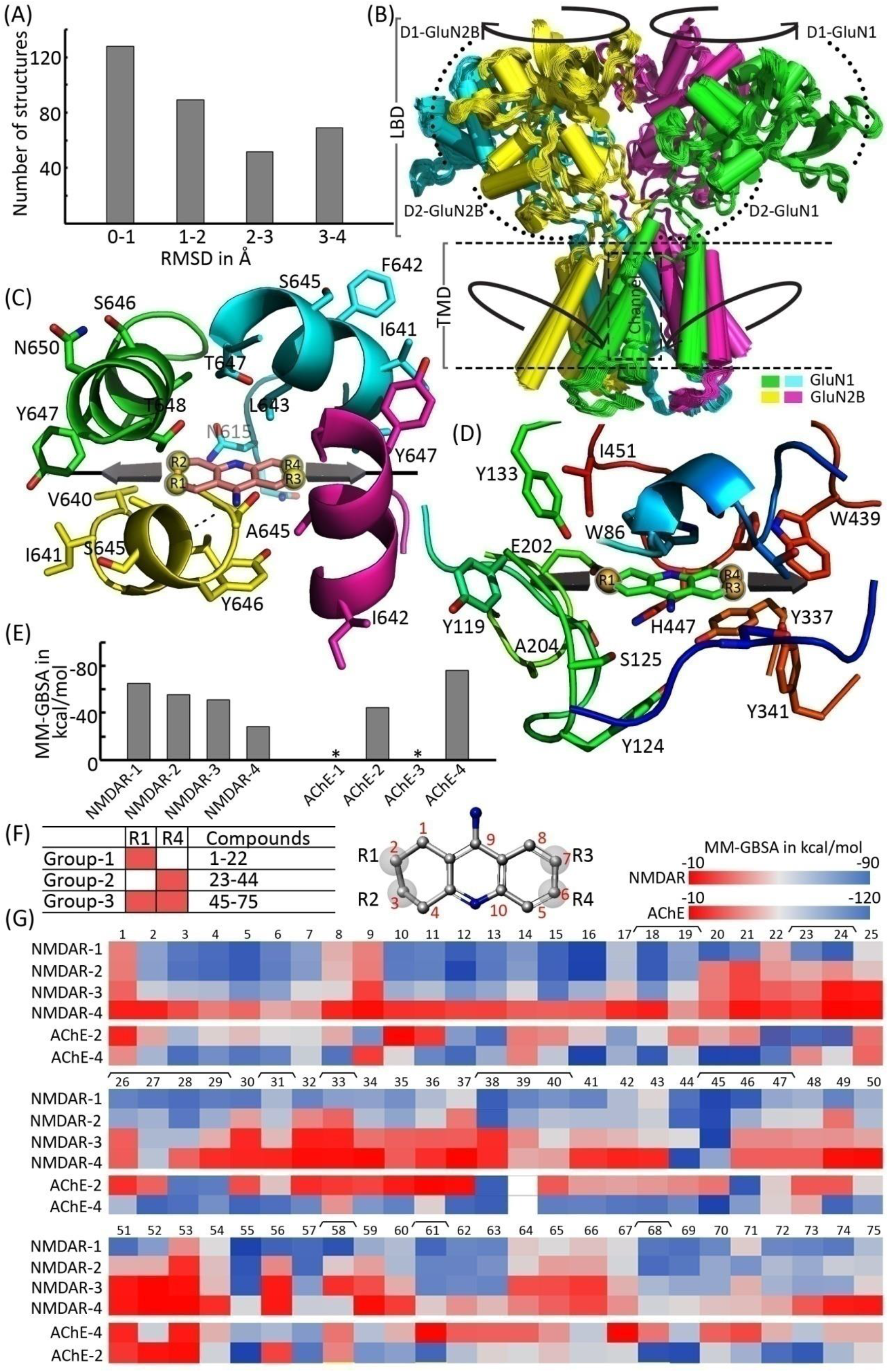
Modeling of h-NMDAR and docking of tacrine derived MTDLs against NMDAR and AChE. **(A)** RMSD based clustering of output structures obtained from the coarse-grained modeling of NMDAR. **(B)** Structure of modeled heteromeric h-NMDAR composed of two subunits each of GluN1 (shown in green and cyan color) and GluN2B (shown in yellow and magenta color). Two key motions featuring shearing/twisting for the LBD (indicated by arrows pointing outwards) and TMD (indicated by arrows pointing inwards) are also shown. **(C)** Binding mode of tacrine (salmon stick) in the channel vestibule of the modeled h-NMDAR. The possible sites available for modifications on tacrine are also denoted as R1 to R4. **(D)** Binding mode of tacrine (green stick) in the active site of human AChE. **(E)** Binding energies (MM-GBSA in kcal/mol) obtained for tacrine from the ensemble docking, where * indicates absence of a biologically significant docked pose. Candidate structures of NMDAR and human AChE used for ensemble docking are represented as NMDAR-1 to NMDAR-4 and AChE-1 to AChE-4 respectively. **(F)** Three-dimensional structure of tacrine marked with available sites for modifications (R1-R4) and number of designed MTDLs that belong to different groups based on the substitutions are also shown. **(G)** Heatmap of the binding energies (MM-GBSA in kcal/mol) obtained for designed MTDLs after ensemble docking. Ligands that are synthesized in the current studies (MTDLs-18, 19, 23, 24, 26-29, 31, 33, 38-40, 45-47, 58, 61 and 68) are marked.

The NMA and ENM simulations predicted two key motions featuring shearing/twisting for the ligand binding domains (LBDs) which is in concert with the transmembrane domain (TMD) (Fig. 1B). It was observed that D1 and D2 lobes of GluN2B move in opposite directions when compared to the motion of D1 and D2 lobes of GluN1, triggering an inward movement for the TMD. Our results are similar to the previously reported structural dynamics of NMDAR [61]. Since the modeled h-NMDAR was in open conformation, it is assumed that, the observed structural dynamics in the coarse-grained modeling might be representing partially or fully closed conformations. In our coarse-grained modeling, it was also observed that the structural changes were transmitting from LBD to TMD through linkers that connect the LBD and three transmembrane helices M1, M3, and M4 at TMD. The surface area of the channel in the candidate structures (denoted as NMDAR-1 to NMDAR-4) were found to be 1269, 832, 801 and 543 Å^2^ respectively. Hence, it is assumed that the conversion from NMDAR-1 to NMDAR-4 might be representing the transformation from open (NMDAR-1) to partially closed (NMDAR-2/3) and to fully closed conformations (NMDAR-4).

#### Binding studies of tacrine towards NMDAR revealed that it binds in the MK-801 binding pocket

Further to understand the binding mode of tacrine, we performed ensemble docking against these four candidate structures. The binding energies of tacrine towards NMDAR-1 to NMDAR-4 are −68, −57, −51 and −38 kcal/mol and these binding energies are directly proportional to the size of the binding site. We selected the best scoring pose of tacrine and further analyzed the atomic interactions. In the most energetically favored binding mode of tacrine, it was observed that tacrine resides within the same pocket where MK-801 binds. Tacrine was oriented in the channel vestibule in such a way that it can form a hydrogen bond with the backbone atom of L643 of GluN2B (Fig. 1C). Other residues in the M3 helices of GluN1 and GluN2B chains were also contributing to the stability of binding.

#### The conformation of Y337 act as a key determinant in case of tacrine binding to AChE

Utilizing the ensemble docking approach, the binding energies of tacrine towards the selected AChE structures were determined. Tacrine was unable to bind to the apo (AChE-1) and donepezil bound (AChE-3) structures of AChE, due to the steric clashes with Y337. Since the side chain orientation of Y337 was favorable in huperzine bound (AChE-2) and 9-aminoacirdine bound (AChE-4) structures, tacrine was able to bind to the active site gorge. Binding energies of tacrine towards AChE-2 and AChE-4 were −42 and −79 kcal/mol respectively. It has been reported that the side chain of Y337 acts as a swinging gate and plays an important role in recognizing different ligands [62]. In the lowest energy pose, tetrahydroacridine moiety of tacrine is sandwiched between W86 and Y337 and thus favored face to face π-π stacking interactions. Similarly, W439 is involved in the edge to face stacking interactions with tacrine. Additionally, a hydrogen bond was observed between backbone O atom of the H447 and N atom of tacrine (Fig. 1D). Binding energies of tacrine towards NMDAR-1 to 4 and AChE-1 to 4 are denoted in Fig. 1E.

#### Structure based design provided several tacrine derived MTDLs

Our critical investigation of the binding modes of tacrine on NMDAR and AChE revealed additional rooms for modifications. We were interested in the four potential pharmacophoric regions in tacrine, represented as R1 to R4 for the designing of potential MTDLs, where R1, R2 are located in the cyclohexane ring and R3, R4 are located in the aromatic ring of tacrine (Fig. 1F). In the case of NMDAR, we found that three hydrogen bond donors (backbone nitrogen atoms of V640, I641 and A644 of GluN2B) and two hydrogen bond acceptors (backbone oxygen atoms of A639 and V640 of GluN2B) are located within 6 Å distance from the R1 position. Hence, we assumed that the substitution of hydrogen bond donors or acceptors at R1 position may favor hydrogen bonding and thus improve the binding affinity towards NMDAR. We then investigated whether these substitutions would also favor the binding towards AChE. In our previous studies, we demonstrated that the substitution at R1 position significantly improved the binding affinities of novel tacrine derivatives [48]. Although, the phenylethylacetamide and phenylpropyl acetamide in compounds 6b, 6c, 6e, 6f, 6h, 6i and 6j maintained AChE inhibitory activities [48], we did not include such bulky substitution in the current study due to the possible steric clashes with residues in the M3 helices of both GluN2B and GluN1.

We then investigated the possibilities of R2 substitutions. In the case of NMDAR, we found only two hydrogen bond donors (hydroxyl group at the side of chain of T648 and backbone N atom of A645 at GluN1) at a distance of 5 Å from the R2 position. In the case of AChE, we found that the R2 position is not favorable for substitution due to the possible steric clashes with the side chain of E202 and backbone N atoms of G120, G121. All these residues were located within 3.5 Å from R2 position. Hence, we omitted R2 position in the current studies from further substitutions.

Our next step was to investigate the possibilities of R3 and R4 substitutions. In our previous studies, we successfully demonstrated that substitutions of bromine and aromatic moieties such as methyl pyrazole and pyrimidine at R4 position maintained the AChE inhibitory activity [48]. In the case of NMDAR, an aromatic cage constituted by the following residues, F548, F554, W563, Y647 (from GluN1) and F637 (from GluN2B) is situated at a distance 10 Å from both R3 and R4. At the same time in AChE, any bulky aromatic substitution at R3 position may make steric clashes with Y341. Based on this observation we decided to eliminate the R3 position for further modifications in the current studies. Moreover, we also found that any bulky aromatic substitutions at R4 position may introduce stacking interactions with W439 and Y449 of AChE. Based on all these observations we designed 75 ligands and grouped them into three different groups where group-1 consists of ligands that are having substitutions only at R1, group-2 consists of substitution only at R4 and group-3 consists of substitutions at both R1 and R4 (Table S1). We then performed ensemble docking studies for all the ligands against both NMDAR and AChE and binding energies were determined. We found the binding energies vary from −10 to −90 kcal/mol towards NMDAR and −10 to −120 kcal/mol towards AChE (Fig. 1G).

#### *In silico* binding affinity guided the selection of tacrine derived MTDLs for synthesis

Though we designed 75 compounds based on the pharmacophoric features derived from tacrine binding, we synthesized only 19 tacrine derived MTDLs for further evaluations. The compounds are named according to our patent application (IPO-201841015699). Corresponding numbers of these compounds in the *in silico* designs are shown in Table S1. *In silico* based ADME properties and binding affinities determined in the current study encouraged us to choose compounds 16 and 17 of group-1, 201, 203-206, 208-212 and 214 of group-2 and 5, 8, 10, 13, 14 and 107 of group-3 for synthesis and further evaluations. No structural alert was found when they were screened for Pan assay interference of compounds (PAINS) using SwissADME tool. The physico chemical properties predicted using the QikProp module for the selected MTDLs are listed in Table S2. The predictions indicate that all compounds have general drug like properties except 206 and 211. The LogP values of these two compounds were slightly higher, but it has already been reported that CNS active ligands can have higher LogP (more lipophilic) values compared to general therapeutics [63, 64] and hence they were included in further studies.

#### Synthesis of selected tacrine derived MTDLs

All the selected MTDLs were synthesized by the reaction procedures described in Schemes 1 and 2 and the final products were characterized by ^1^H NMR and LCMS. The 6-bromo tacrine (3) was synthesized by condensation of 4-bromo-2-amino benzonitrile (1) and cyclohexanone (2) in presence of borane trifluoro etherate (BF_3_.Et_2_O) under reflux for 4 hours. The palladium-mediated Suzuki-Miyaura cross coupling reaction of compound 3 with corresponding boronic acids or boronic esters (4a-4m described in Experimental section) resulted in the title compounds 201 and 203-214. Similar procedure was adopted to get the compounds 05, 16, 10 and 107. Compounds 05 and 10 were further treated with aqueous methyl amine to synthesize compounds 13 and 14 respectively. The compounds 17 and 08 were synthesized from 16 and 05 respectively by reacting with hydrazine hydride.

**Scheme 1:**
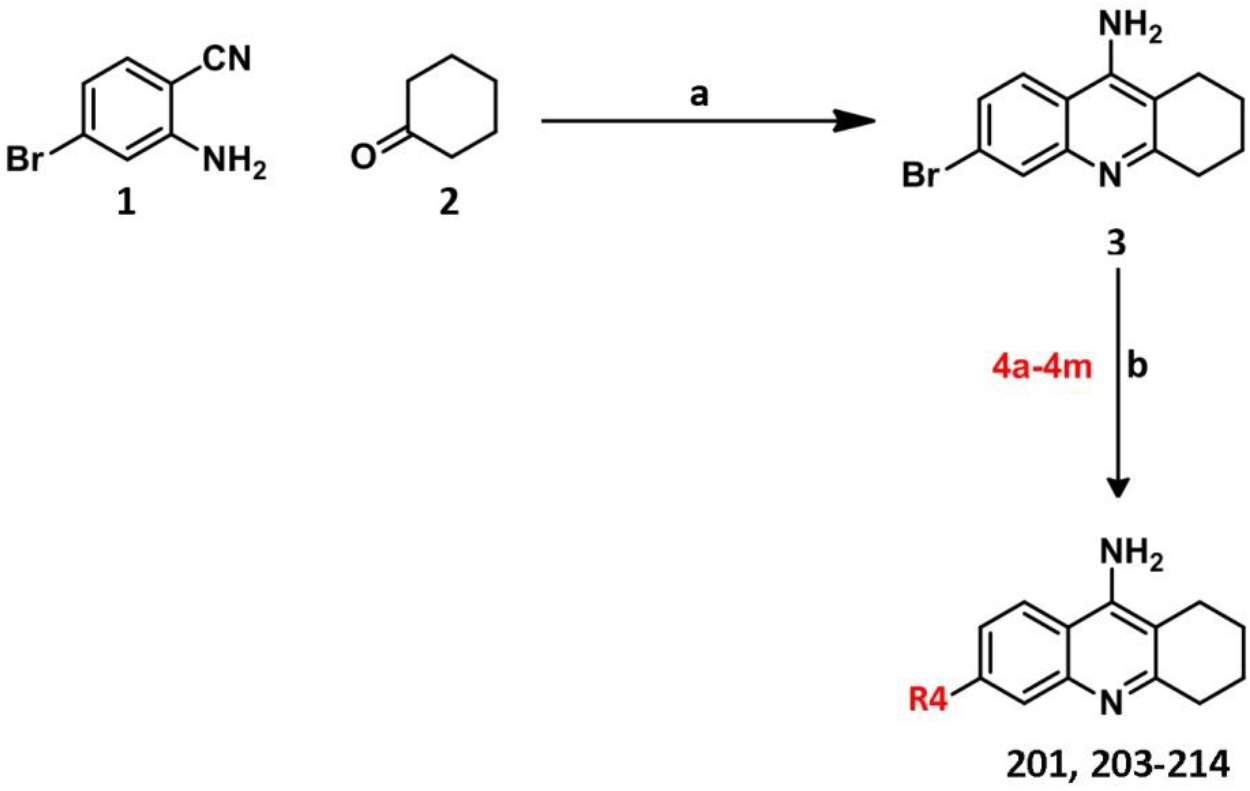
Synthesis scheme for R4-substituted tacrine derivatives

#### Reagents and conditions

(a) BF_3_.Et_2_O, toluene, 110°C, 4 h (b) compounds 4a-4m, Na_2_CO_3_, Pd(PPh_3_)_4_, 1,4-dioxane and water (9:1), 110°C, 2 h, seal tube

**Scheme 2:**
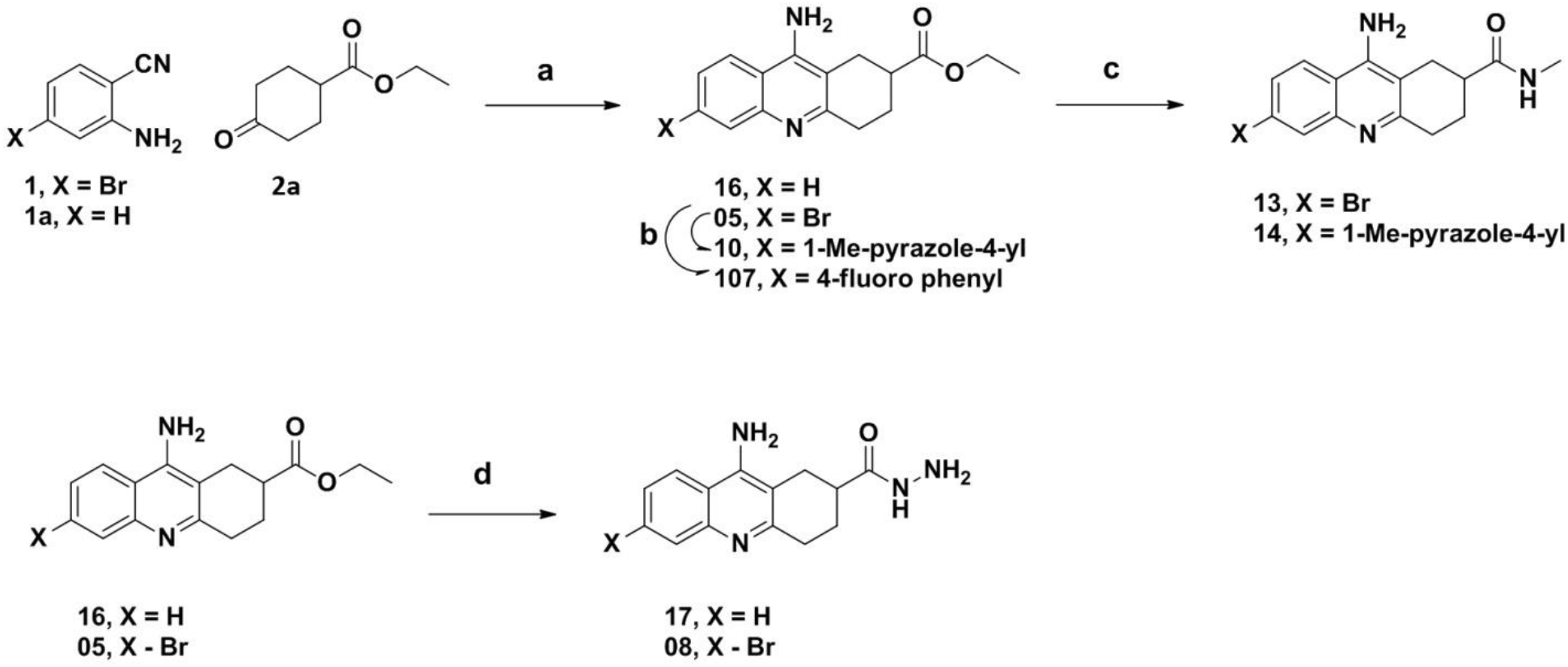
Synthesis scheme for R1 and R4-substituted tacrine derivatives

#### Reagents and conditions

(a) BF_3_.Et_2_O, toluene, 110°C, 4 h (b) compound **4a** for **10** and **4h** for 107, Na_2_CO_3_, Pd(PPh_3_)_4_, 1,4-dioxane and water (9:1), 110°C, 2 h, seal tube; (c) aq.MeNH_2_, 60°C, 4 h; (d) NH_2_NH_2_.H_2_O, 60°C, 2 h

### *In vitro* studies

#### Tacrine derived MTDLs inhibit cholinesterases (ChE)

Ellman’s method based ChE assay revealed varying degrees of inhibitory activities for MTDLs on ChEs. Our *IC*_*50*_ evaluation of tacrine against BChE (*IC*_*50*_=14.26±1.07 nM) and AChE (*IC*_*50*_=94.69±4.88 nM) confirmed the selectivity of tacrine towards BChE over AChE as reported in the literature [65]. The *IC*_*50*_ values of all MTDLs against AChE were in the nM range and towards BChE were in the µM range (Fig. 2A and 2B). Hence, we concluded that these MTDLs were more selective towards AChE than BChE.

**Figure 2:**
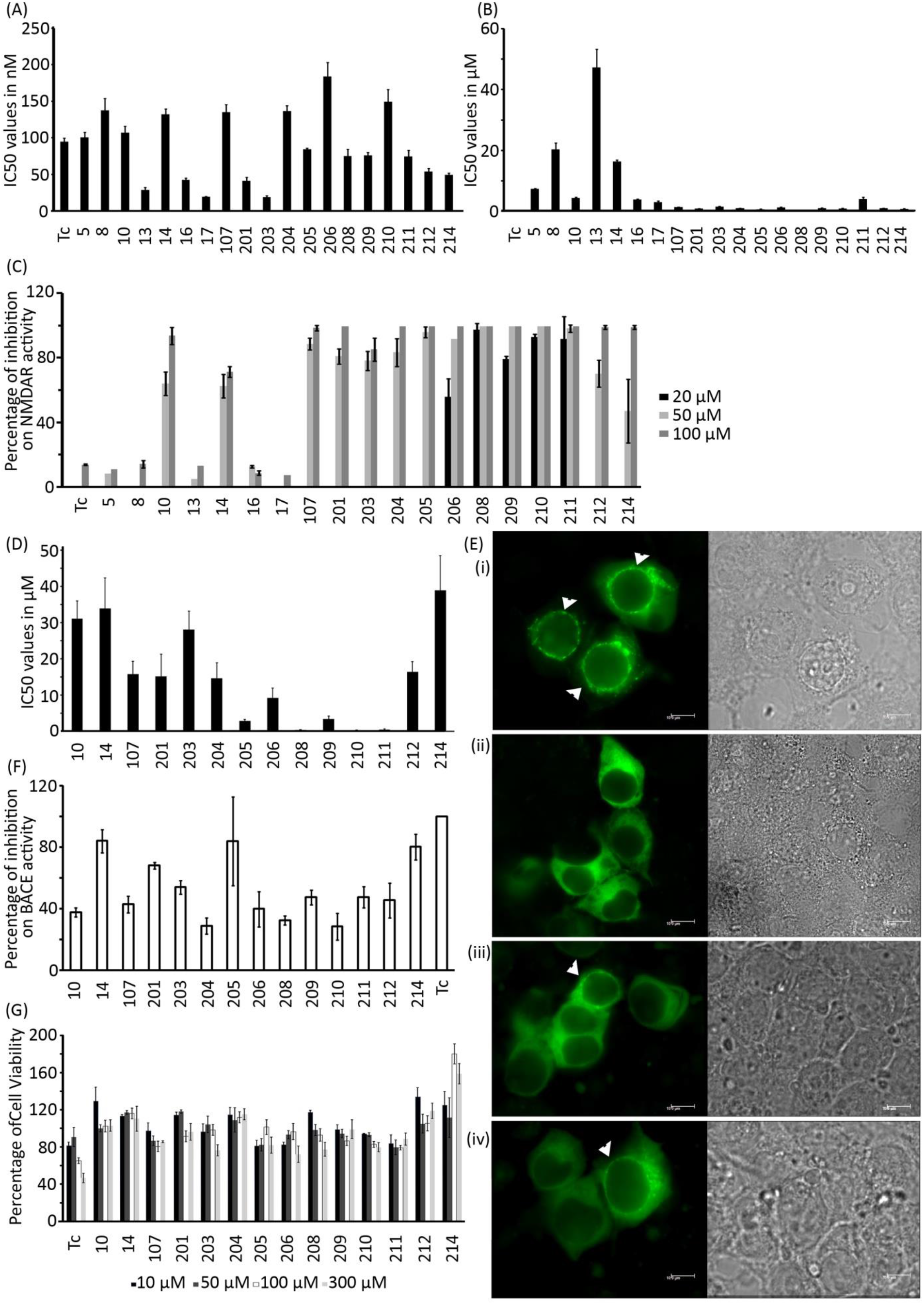
*In vitro* inhibitory profile of MTDLs against different targets. *IC*_*50*_ values (represented as mean±SD, n=3) determined for the tacrine derived MTDLs towards AChE and BChE are shown in **(A)** and **(B)** respectively. **(C)** Initial screening of MTDLs and tacrine (Tc) for inhibition on NMDAR activity. The values presented are mean±SD (n =3). **(D)** *IC*_*50*_ values (represented as mean±SD, n=3) determined for tacrine derived MTDLs towards NMDAR are shown. **(E)**Representative images of HEK-293 cells expressing GFP-α-CaMKII, GluN1 and GluN2B. (i) Punctate appearance in the presence of NMDAR agonists (glutamate and glycine) and calcium. (ii) Absence of punctate appearance in the presence of agonists but in the absence of Ca^2+^; (iii) Reduction in the number of punctate cells when treated with NMDAR agonists, Ca^2+^ and 211 at 5μM; (iv) Reduction in the number of punctate cells when treated with NMDAR agonists, Ca^2+^ and 208 at 5μM. Right panel shows the corresponding bright field images. **(F**) Percentage of residual BACE activity (represented as mean±SD, n=3) in the presence of 50 μM tacrine (Tc) or tacrine derived MTDLs. **(G)** Cytotoxicity of the compounds on HepG2 cell line (represented as mean ±SD, n=3) after exposure for 24 hours is shown.

Our detailed analysis of the *IC*_*50*_ values revealed that 11 (Compound numbers-13, 16, 17, 201, 203, 205, 208, 209, 211, 212 and 214) out of 19 compounds exhibited higher inhibition towards AChE than tacrine. Among these compounds, compound 203, having a pyrimidine ring connected to the R4 position hasan excellent anti AChE activity (*IC*_*50*_=18.53±2.09 nM). Substitutions of methyl pyrazole (compound 201, *IC*_*50*_=40.89±4.82 nM) and fluoro benzene (compound 209, *IC*_*50*_=76±3.79 nM) at R4 position of tacrine were found to have more inhibitory potency compared to compounds with substitutions at R1 position (compounds 10: *IC*_*50*_=107.17±8.68 nM, 14: *IC*_*50*_=131.92±7.72 nM and 107: *IC*_*50*_=135.11±10.25 nM). Simple halogen substitutions at R4 position along with substitutions at R1 position (compounds 5: *IC*_*50*_=100.4±7.2 nM and compound 8: *IC*_*50*_=137.7±16.1 nM) exhibited less inhibitory activity when compared to substitutions at R1 alone (compounds 16: *IC*_*50*_=42.2±2.7 nM and 17: *IC*_*50*_=19.27±0.47 nM). The study also revealed that compounds with simple aromatic ring at R4 position (compounds 201, 203, 212 and 214) are preferred over the compounds with substituted aromatic rings (compounds 205, 206, 208, 209, 210 and 211).

#### Improved antagonistic potential of tacrine derived MTDLs towards NMDAR

The effect of MTDLs on NMDAR activity was evaluated using a cell-based assay system, in which the appearance of fluorescent punctate pattern occurred during the activation of NMDAR by its agonists and calcium (Ca^2+^) (Fig. 2E (i)) [66]. The punctate appearance was not observed when NMDAR was treated with its agonists in the absence of Ca^2+^ (Fig. 2E (ii)), which clearly indicated that the assay system is dependent on Ca^2+^ influx through NMDARs. Activationof NMDAR in presence of the MTDLs resulted in the reduction of the number of cells with punctate pattern which is positively correlated to the inhibition of Ca^2+^ influx through NMDAR. Initially, two different concentrations (50 and 100 μM) of the compounds were used for testing their effect on NMDAR activity.

All MTDLs except tacrine and compounds 5, 8, 13, 16 and 17 exhibited >60% inhibition of NMDAR activity (Fig. 2C). Almost 100% inhibition was observed for the compounds 201, 204, 205, 206, 208, 209, 210 and 211 at 100 μM and for compounds 208, 209, 210 and 211 this happened at 50 μM itself. Further, their inhibitory potency at 20 μM was also tested in order to determine the concentration range for inhibition. From the initial screening studies, it was found that MTDLs belonging to group-2 having substitutions only at R4 position (i.e. compounds 201, 203, 204, 205, 206, 208, 209, 210, 211, 212 and 214) were effective in antagonizing NMDAR activity while the compounds belonging to group-1 where the substitution was only at R1 position (compounds 16 and 17) could not effectively block NMDAR activity. MTDLs that belong to group-3 with dual substitutions i.e. aromatic substitutions at R4 and substitutions at R1 positions (compounds-10, 14 and 107) inhibited NMDAR activity better when compared to simple halogen substitutions at R4 position (compounds 5, 8 and 13). Hence, we inferred that aromatic substitutions at R4 position alone might be sufficient to antagonize NMDAR activity, which is corroborated by *IC*_*50*_ determinations. It was clear from the measured *IC*_*50*_ values (Fig. 2D) that the inhibitory potency of some MTDLs (208, *IC*_*50*_=0.31±0.09 μM and 210, *IC*_*50*_=0.27±0.05 μM) were almost 600 fold higher when compared to the reported value for tacrine (*IC*_*50*_=193±33 μM) [37]. About 4 fold increase in potency was observed for compound 209 (*IC*_*50*_=3.36±0.86 μM) when compared to 107 (*IC*_*50*_=15.81±3.44 μM) and the potency was further increased upto 10 fold by halogen substitution at ortho position of phenyl ring at R4 position of tacrine (compound 208). Representative images showing NMDAR activities in presence of agonists and in the presence of compounds 211 and 208 are shown in Fig. 2E (iii and iv).

#### MTDLs do not interfere with the interaction between GluN2B and α-CaMKII

In the NMDAR assay, activity is detected based on the formation of punctae as a result of Ca^2+^-dependent protein-protein interaction between CaMKII and GluN2B subunit of NMDAR. Any agent which can block the interaction between these two proteins may reduce or completely inhibit the appearance of punctae. This reduction in punctae would not be due to the blocking of NMDAR channel activity. To test whether the compounds have any effect on the interaction of GluN2B and α-CaMKII, an experiment was carried out using HEK-293 cells stably expressing GFP-α-CaMKII and GluN2B sequence [66]. Formation of perinuclear punctae by treatment with ionomycin and Ca^2+^, is indicative of interaction between CaMKII and GluN2B. No detectable reduction in the punctae formation in presence of the MTDLs confirmed that the compounds do not have any effect on the interaction of GluN2B and α-CaMKII (Fig. S1 and S2).

#### β-site amyloid precursor protein cleaving enzyme (BACE-1) activity reduced in presence of tacrine derived MTDLs

Since BACE-1 is a rate limiting enzyme in the production of amyloid beta peptide that contributes to AD pathology, we checked the inhibitory effect of tacrine derived MTDLs on BACE-1 activity. Five compounds namely 10, 204, 206, 208 and 210 efficiently inhibited (>60%) BACE-1 enzyme activity at 50 μM (Fig.2F).

#### MTDLs are less hepatocytotoxic than tacrine

Since hepatotoxicity is one of the serious concerns with new tacrine derivatives, we checked the cytotoxic nature of all derivatives on liver carcinoma cell line, HepG2 using 3-(4,5-dimethylthiazol-2-yl)-2,5-diphenyltetrazolium bromide (MTT) assay. HepG2 cell line is one of the widely used *in vitro* models to study hepatotoxicity of chemicals [67]. As seen in Fig. 2G, tacrine was safe only up to 50 μM and the cell viability decreased from 100 μM onwards. At 300 μM, almost 50% reduction in cell viability was observed. Almost all tacrine derived MTDLs tested were found to be less toxic than tacrine (approximately >80% cell viability at 300μM). Compounds such as 14, 201, 204, 209, 212 and 214 were non-toxic even at a concentration of 300 μM.

### Neuroprotective properties of tacrine derivatives

#### Establishment of glutamate toxicity model in rat primary cortical neurons

The neuroprotective properties of selected MTDLs were evaluated against glutamate induced excitotoxicity in primary cortical neurons prepared from the brain of embryonic rats. Cortical neurons were maintained in culture and excitotoxicity experiments were performed on the 9^th^ day of culture (DIV9). Previous studies demonstrated that neurons at DIV9 are appropriate to study glutamate toxicity as the cells are mature enough in terms of receptor expression and glutamate sensitivity [68–71]. An *in vitro* excitotoxicity model was established by treating cortical neurons with 100 μM glutamate for 1 hour followed by analysis at 24 hours post treatment. The extent of excitotoxic cell death was quantified biochemically by measuring the activity of glucose-6-phosphate dehydrogenase (G6PD) released into the medium (Fig. 3A). As expected, treatment with glutamate led to increased cell death which was prevented by the NMDAR inhibitor, MK-801. The results substantiated that cell death in this model is primarily mediated by NMDAR activation. Cell death was assessed by imaging after immunostaining for the neuronal marker protein, microtubule associated protein 2 (MAP2) and also by observing chromatin condensation as detected by nuclear staining with DAPI (Fig. 3B). Immunocytochemical staining of neurons for MAP2, predominantly localized in dendrites [72] has been used as a marker to study neuronal integrity [73] and thus neuronal survival [74]. DAPI and immunostaining showed a dramatic decrease in the viable and MAP2 positive neurons upon glutamate treatment. Staining experiments also showed that excitotoxicity induced by glutamate is prevented in the presence of MK-801. The percentage of viable cells and percentage of MAP2 positive neurons in the presence and absence of glutamate are shown in Fig. 3C. Glutamate mediated excitotoxicity was also checked using MTT assay 24 hours after treating cultures with glutamate for 3 hours. Compared to the glutamate free control, glutamate treatment showed a significant reduction in neuronal viability (Fig. 3D).

**Figure 3:**
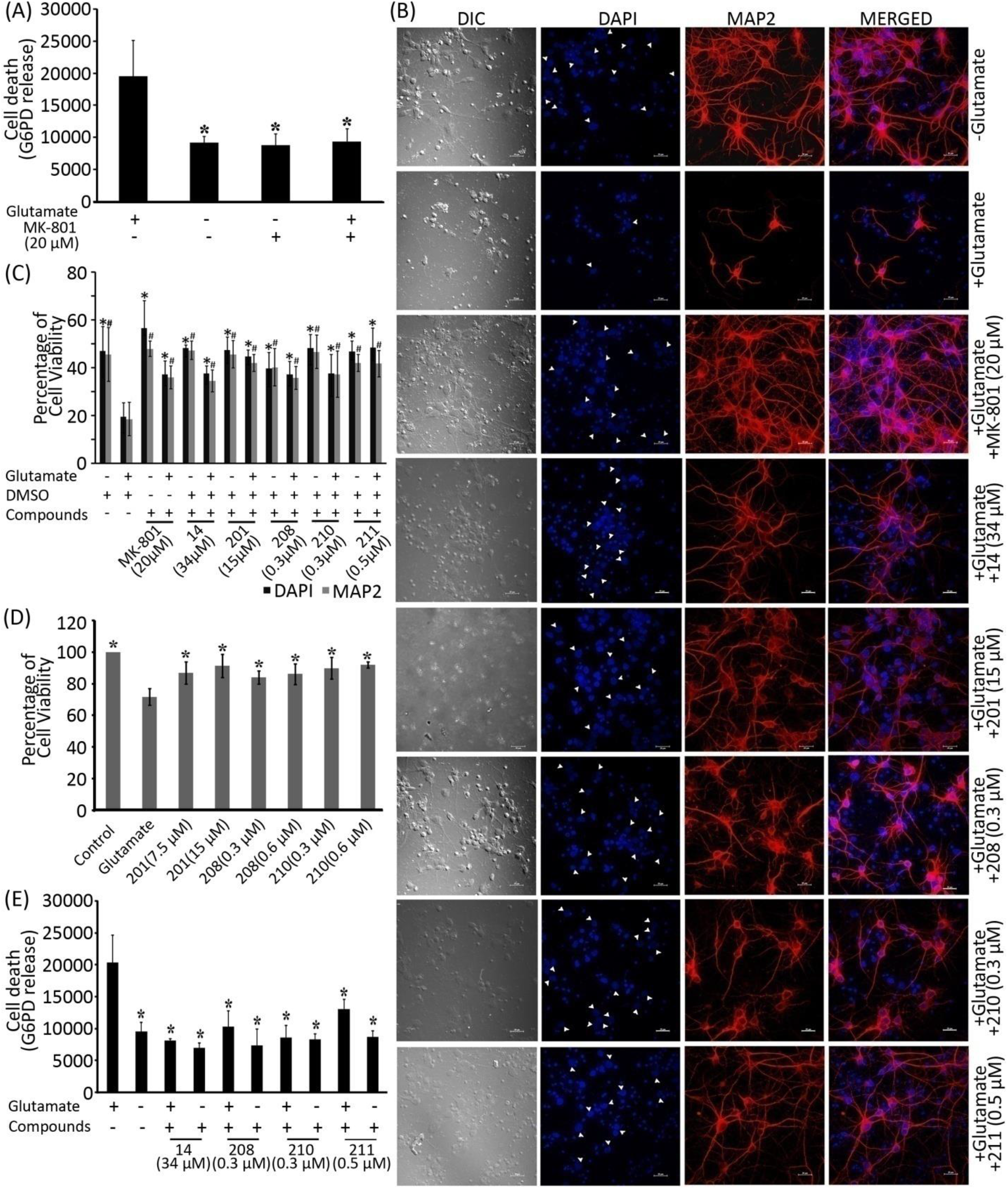
Neuroprotective action of tacrine derived MTDLs on glutamate induced excitotoxicity in primary cortical neurons at DIV9 treated with 100 μM glutamate and 10 μM glycine. **(A)** The extent of G6PD released as a part of excitotoxic cell death in the presence and absence of glutamate (represented as mean±SD, n=3), * indicates p value <0.05 compared to glutamate treatment. **(B)** Representative images of primary cortical neurons immunostained for MAP2 (red) and counterstained with DAPI (blue) after treatment with 100 μM glutamate and 10 μM glycine in the absence or presence of MK-801 or selected tacrine derived MTDLs (14, 201, 208, 210 and 211). Live neurons are shown with arrow heads. **(C)** The quantification of glutamate-induced cell death detected by DAPI staining and MAP2 immuno-reactivity in cortical neurons. The bar diagrams represent the percentage of viable and MAP2 positive cells (represented as mean±SD, n=3). *and # indicates p value <0.05 compared to glutamate treatment for the viable cell count and MAP2 positive neurons respectively. **(D)** Glutamate induced neuronal death in the absence or presence of selected MTDLs (201, 208 and 210) measured using MTT assay (represented as mean±SD, n=3). * indicates p value <0.05 compared to glutamate treatment. **(E)** Glutamate induced death of neurons in the absence or presence of selected MTDLs (14, 208, 210 and 211) measured using G6PD release assay (represented as mean±SD, n=3), DMSO was present in all samples. * indicates p<0.05 compared to glutamate treatment.

#### Treatment with tacrine derived MTDLs reduces glutamate induced toxicity

Based on the NMDAR inhibition pattern, our MTDLs can be classified into three categories of inhibitors such as potent (*IC*_*50*_<5 μM), moderate (*IC*_*50*_<20 μM) and less potent (*IC*_*50*_>20 μM) ones. We selected candidates from each category to assess their neuroprotective properties. Totally five of the MTDLs (14, 201, 208, 210 and 211) were taken to assess their effects on glutamate induced excitotoxicity, of which three (208, 210 and 211) are potent blockers of NMDAR. The remaining two, 201 and 14, showed moderate and reduced potency towards NMDAR respectively. Biochemical assays such as G6PD release assay or MTT assay were performed to estimate the neuroprotective effect. All compounds were dissolved in DMSO (< 0.05%) as it has been shown that DMSO does not have any protective effect against glutamate toxicity (Fig. S3). Controls with DMSO were used in all experiments.

Assays for G6PD release show that compounds 14, 208, 210 and 211can protect neurons against glutamate toxicity at concentrations close to their respective *IC*_*50*_ values (Fig. 3E). The reduction in neuronal viability by glutamate treatment was averted in presence of compounds 201, 208 and 210 as evident from MTT assay. In these experiments neurons were treated with compounds along with glutamate after pre-treatment with compounds (Fig. 3D). Both the bio-chemical results clearly illustrate that these compounds can protect cortical neurons from glutamate induced toxicity.

#### Treatment with tacrine derived MTDLs prevents glutamate induced neuronal injury

The protective effect of the compounds on glutamate-induced neuronal death was further studied by nuclear staining using DAPI. Glutamate treatment in presence of compounds 14, 201, 208, 210 and 211 showed a clear decrease in the number of condensed nuclei compared to glutamate treatment in the absence of the compounds (Fig. 3B and 3C). This is an indication of neuroprotection by compounds against excitotoxicity.

#### Treatment with tacrine derived MTDLs prevents loss of neuronal integrity

The effect of MTDLs 14, 201, 208, 210 and 211 on glutamate induced loss of neuronal integrity was studied by immunocytochemical staining for MAP2. Decrease in MAP2 positive cells and loss of neuronal architecture caused by glutamate treatment were prevented in presence of the MTDLs which pointed towards the protective role of these derivatives (Fig. 3B and 3C). No major changes were induced in neurons when treated with the compounds in the absence of glutamate (Fig. S3).

### *In vivo* studies

#### Tacrine derived MTDLs ameliorated mono sodium glutamate (MSG) induced cognitive impairment

MSG has been shown to induce a series of behavioral disorders and brain lesions in various experiments by stimulating glutamate receptors [75–79]. Based on this information, MSG was used to build chronic excitotoxic conditions in a rat model to study the neuronal damage induced learning impairment using the MWM test. MSG (2g/Kg body weight) was given intraperitoneally (IP) for 15 days. We selected tacrine, compounds 201 and 208 to assess whether these compounds can ameliorate MSG induced cognitive impairment. We administered these compounds by IP injection simultaneous to MSG treatment as described in the experimental section. The test included 3 days of training in the Morris water maze. The mean escape latency values of all the groups on each day are shown in Fig. S4. Compared to saline treated group, MSG treated animals showed significant increase in latency to reach the platform on the 3^rd^day of the trials (Fig. 4A) indicating impaired learning. The compounds 201 and 208 when administered along with MSG caused significant improvement in escape latency (Fig. 4A) showing that the compounds were able to ameliorate the impaired performance caused by administration of MSG. Tacrine with MSG did not exhibit any significant improvement in learning ability when compared to MSG alone.

**Figure 4:**
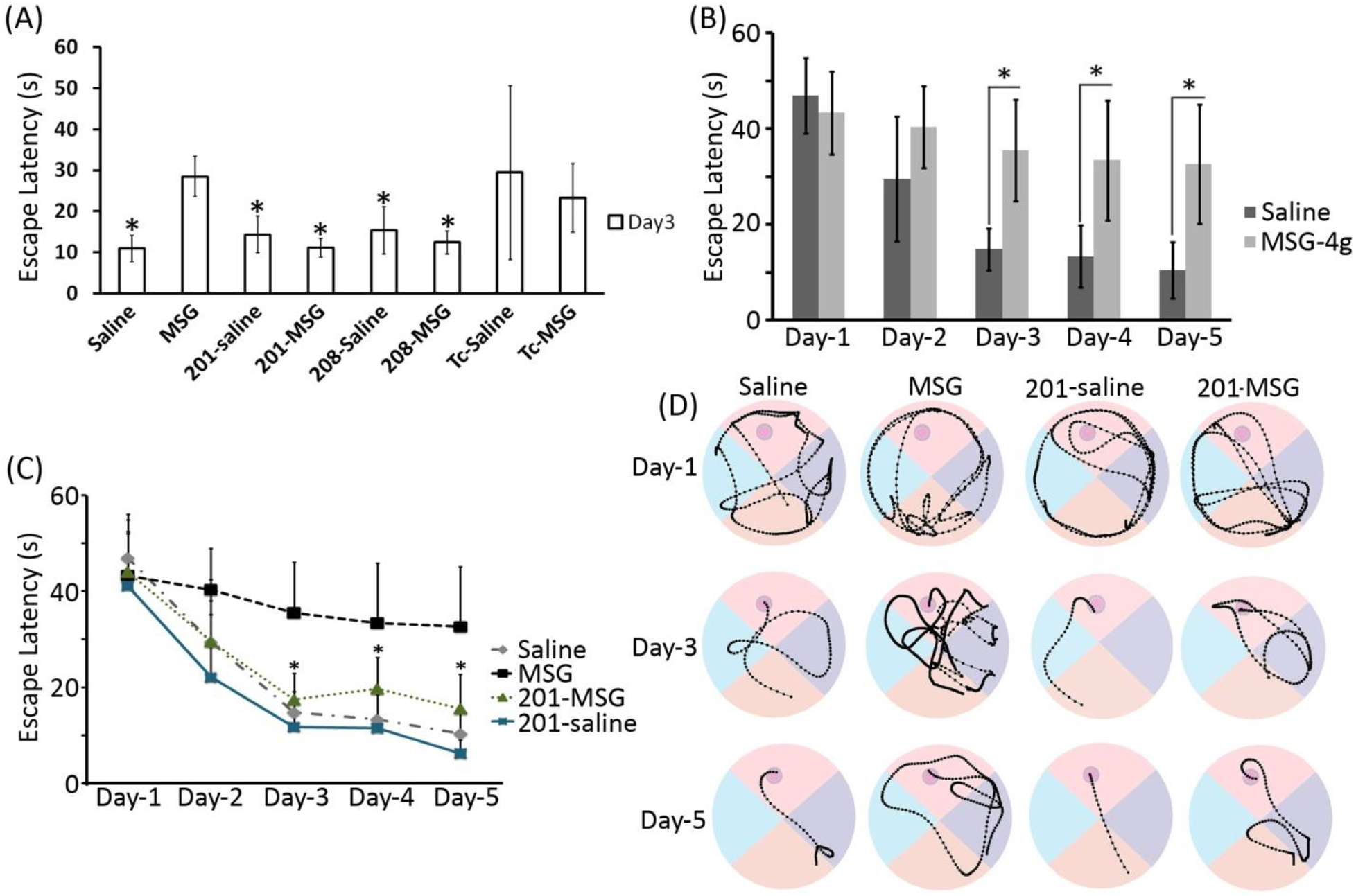
The *in vivo* effect of selected MTDLs (201 and 208) assessed using MWM behavioral test. Animals were subjected to treatment with saline or MSG along with the MTDL or vehicle. **(A)** Escape latencies of animals after the indicated treatments (represented as mean±SD, n=3) to reach the platform on 3^rd^ day of trial, *p value <0.05 compared to treatment with MSG alone (2 g/Kg body weight). **(B)** Effect of higher dose of MSG (4 g/Kg bodyweight) on learning and memory task compared to saline control. Escape latencies of the animals (represented as mean±SD, n=7) from Day1 to Day5 are shown, * indicates p value <0.05 compared to corresponding saline of the same day. **(C)** Effect of compound-201 on cognitive impairment induced by MSG (4 g/Kg bodyweight). Escape latencies from Day1 to Day5 are shown (represented as mean±SD, n=7 for saline, MSG and 201-saline groups and n=6 for 201-MSG group). * indicates p value <0.05 compared to MSG treatment alone. The data for saline and MSG is same as that shown in B. **(D)**Representative trajectories of selected animals in each group in the MWM experiment on days 1, 3 and 5.

We further conducted experiments with the compound 201, to confirm its *in vivo* neuro-protective effect on learning and memory. Even though 201 has shown only moderate affinity towards NMDAR compared to 208, the non-toxic behavior of 201 on the HepG2 cell line compared to 208 encouraged us to select 201 for further studies. We used a higher dose of MSG i.e., 4g/Kg body weight to induce excitotoxicity and the MWM test was conducted for 5 days. We found that latency to reach the platform for the MSG-treated group was significantly higher when compared to the control group upto 5 days (Fig. 4B). Treatment of 201 (3 mg/Kg body weight) along with MSG significantly reduced the escape latency compared to MSG alone, signifying that 201 could ameliorate the memory impairment caused even by higher dose of MSG (Fig. 4C). The minor difference between the 201-saline and 201-MSG groups was not statistically significant. The representative trajectories of the trials are given in Fig. 4D.

## Discussion

Since 201 and 208 (differ only in the substitutions at the R4 position, where 201 and 208 have methylpyrazole and fluorobenzene moieties at the R4 position respectively) were promising in all the experiments, pharmacokinetic properties of these MTDLs were predicted using Swiss ADME and were compared with that of tacrine. The pharmacokinetic and pharmacodynamic properties of tacrine are already known [80] and served as a positive control in our molecular modeling studies.

The bioavailability radar plot enabled a first glance at the drug likeness of the compounds. Six physicochemical properties are considered: lipophilicity, size, polarity, solubility, flexibility and saturation. The physicochemical range on each axis is depicted as a colored zone in the radar plot, in which the molecules that fall entirely into the purple area are considered to be drug-like. The physicochemical properties of tacrine (Fig. 5A) and 201 (Fig. 5B) completely fall into the colored zone indicating their drug likeliness. The value of insaturation is slightly exceeding the recommended limit in the case of 208 (Fig. 5C). The tool also predicted that the compounds are blood-brain barrier (BBB) permeable. Since BBB is considered to be the bottleneck in CNS drug discovery, we have also checked the BBB permeability of the compounds by other online tools such ‘online BBB predictor’ and QikProp (Schrödinger). Online BBB predictor was built by applying the support vector machine (SVM) and Ligand Classifier of Adaptively Boosting Ensemble Decision Stumps (LiCABEDS) algorithms [81] and was specially designed to classify compounds based on whether they can cross the BBB (BBB+) or not (BBB−). According to the tool, the threshold value for a compound to be BBB permeable is 0.02. The values predicted for tacrine, 201 and 208 are 0.120, 0.141 and 0.140 respectively indicating that all these compounds are BBB permeable (Fig. 5D). According to QikProp results both 201 and 208 are BBB permeable and CNS active (Table S2).

**Figure 5:**
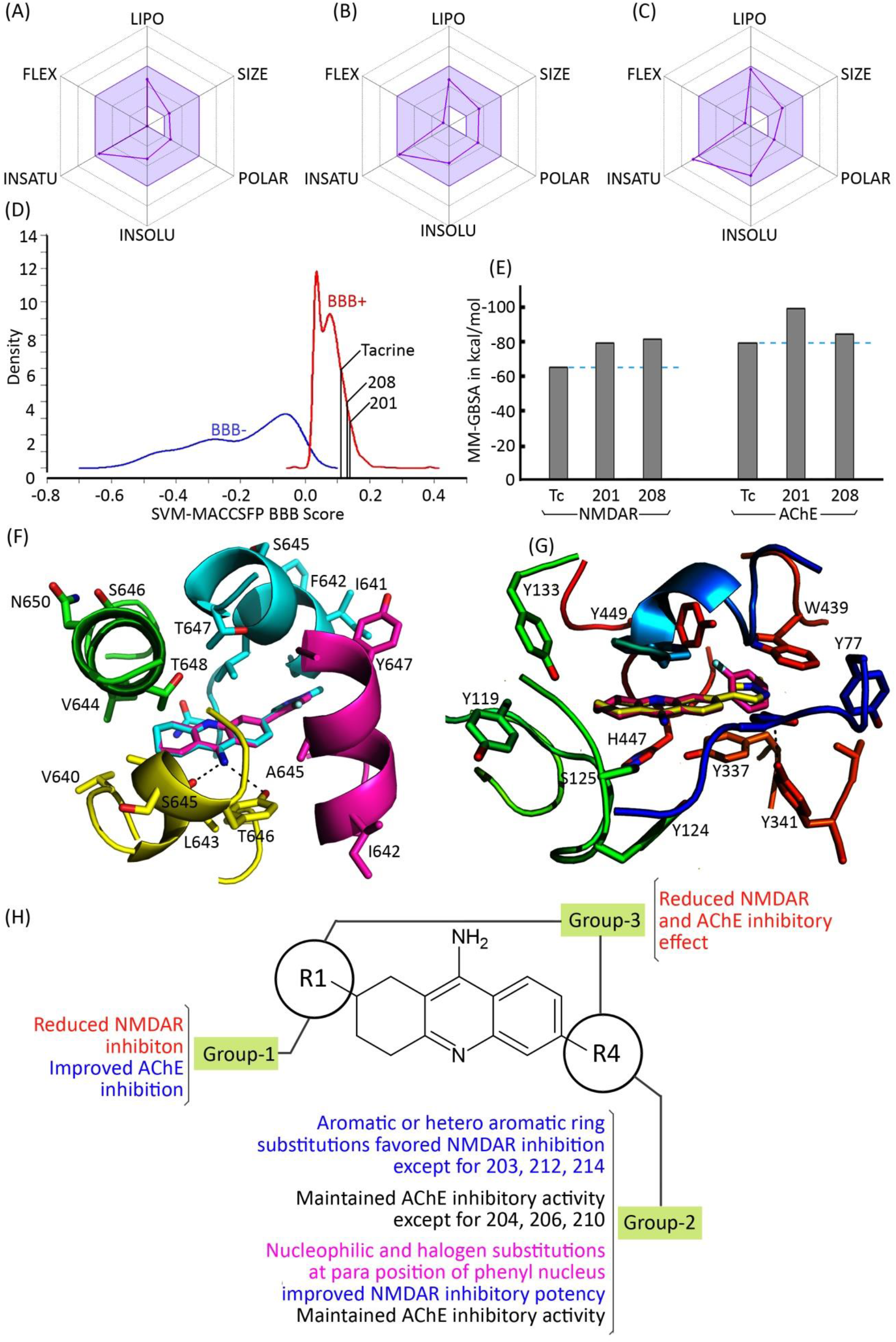
*In silico* ADME properties, binding mode and binding energies determined for tacrine and selected tacrine derived MTDLs 201 and 208. Oral bioavailability radar plots of tacrine **(A)**, 201 **(B)** and 208 **(C)** predicted using SwissADME, in which the colored zone represents the suitable physico-chemical space for oral bioavailability. The optimum range of these properties are as follows: LIPO (Lipophilicity): −7.0<XLOGP3<+5.0; SIZE:150 g/mol<MW<500g/mol; POLAR (Polarity): 20Å^2^<TPSA<130Å^2^; INSOLU (Insolubility): 0<Log S (ESOL)<6; INSATU (Insaturation): 0.25<Fraction Csp3< 1; FLEX (Flexibility): 0< No. of rotatable bonds<9. **(D)** Prediction of blood brain barrier permeability using ‘online BBB predictor’. The values (SVM-MACCSFP BBB score) predicted for tacrine, 201 and 208 are 0.120, 0.141 and 0.140 respectively. Reference range of SVM-MACCSFP BBB score for training set with (marked in red) and without (marked in blue) BBB penetrability are also shown. All compounds are predicted to be BBB+. **(E)** The binding energies calculated for tacrine, 201 and 208 against h-NMDAR and human AChE using Prime MM-GBSA method. **(F)** Binding modes of 201 (Cyan stick) and 208 (Purple stick) in the channel vestibule of modeled h-NMDAR (Cartoon representation). **(G)** Binding modes of 201 (Yellow stick) and 208 (Purple stick) at the active site gorge of human AChE (cartoon representation). Hydrogen bonds are shown as dotted line in both **(F)** and **(G)**. **(H)** Structure vs activity relationship of the synthesized molecules.

Our docking studies revealed that both 201 and 208 bind to AChE (Binding energy (ΔG) for 201 and 208 are −100 and −82 kcal/mol respectively) and NMDAR (ΔG for 201 and 208 are −80 and −82 kcal/mol respectively) much stronger than tacrine (ΔG for tacrine towards AChE and NMDAR −79 and −68 kcal/mol respectively) (Fig. 5E). Both 201 and 208 bind to the active site of AChE and channel vestibule of NMDAR in a similar fashion as tacrine. We found that the tacrine moieties in 201 and 208 were well aligned to tacrine. The substitutions of methyl pyrazole in 201 and fluorophenyl in 208 favoured stacking interactions with Y337 in AChE. Additionally, a hydrogen bond between hydroxyl group of Y341 and N atom of the pyrazole ring of 201 was observed (Fig. 5F). Both of these aromatic moieties improved the hydrophobic contacts with the surrounding residues. Binding modes of 201 and 208 at the channel vestibule of NMDAR are highly similar to each other; the substitutions on the tacrine moiety were oriented between the M3 helices of GluN1 and GluN2B. The interactions of these compounds are largely favored by hydrophobic contacts by the residues in the M3 helices of both GluN1 and GluN2B (Fig. 5G). Since compounds 201 and 208 were promising and belong to the group-2, we compared the *in vitro* inhibitory activities of other molecules in group-2 (Table 1). We found that aromatic or hetero aromatic ring substitutions at R4 position (201, 205, 208, 209, 211 and 212) favoured the inhibitory activity against AChE and NMDAR. Also, halogen substitutions in the phenyl ring at R4 position favoured inhibitory potency towards AChE and NMDAR (208 and 209). However, group-3 compounds that have substitutions at R1 and R4 positions (10 and 14) showed reduced inhibitory activity towards NMDAR and AChE. Although the group-1 compounds exhibited reduced inhibitory activity towards NMDAR, their binding towards AChE was promising. A simple schematic representation of the structure activity relationships (SAR) inferred based on the available data is shown in Fig. 5H.

**Table 1:** Results of AChE, BChE, NMDAR and BACE-1 inhibition studies of the tacrine derived MTDLs, n.d indicates not determined. Data is the same as that shown in Figure2.

| *IC*<sub>50</sub> values (mean±SD) |  |  |  |  |  |
| --- | --- | --- | --- | --- | --- |
| Sl No | Compound numbers | AChE (nM) | BChE (μM) | NMDAR (μM) | % of residual BACE-1 activity (at 50 μM) |
| 1 | 5 | 100.4±7.2 | 7.38±0.21 | n.d | n.d |
| 2 | 8 | 137.7±16.1 | 20.39±2.11 | n.d | n.d |
| 3 | 10 | 107.17±8.7 | 4.39±0.28 | 31.12±4.8 | 37.76±2.89 |
| 4 | 13 | 28.59±3.59 | 47.29±5.98 | n.d | n.d |
| 5 | 14 | 131.92±7.7 | 16.39±0.54 | 33.86±8.41 | 84.11±7.59 |
| 6 | 16 | 42.2±2.7 | 3.82±0.25 | n.d | n.d |
| 7 | 17 | 19.27±0.47 | 2.99±0.40 | n.d | n.d |
| 8 | 107 | 135.11±10.3 | 1.35±0.09 | 15.81±3.44 | 42.97±5.40 |
| 9 | 201 | 40.89±4.82 | 0.85±0.01 | 15.17±6.14 | 68.09±2.09 |
| 10 | 203 | 18.53±2.09 | 1.50±0.20 | 28.1±5.1 | 54.16±4.43 |
| 11 | 204 | 136.70±7.35 | 0.96±0.07 | 14.59±4.32 | 28.95±5.31 |
| 12 | 205 | 84.17±1.80 | 0.70±0.08 | 2.84±0.46 | 84.03±28.93 |
| 13 | 206 | 184.09±19.23 | 1.21±0.16 | 9.19±2.68 | 39.88±11.42 |
| 14 | 208 | 75.07±9.46 | 0.13±0.02 | 0.31±0.09 | 32.53±2.94 |
| 15 | 209 | 76.00±3.79 | 1.03±0.17 | 3.36±0.86 | 47.56±4.76 |
| 16 | 210 | 149.27±16.82 | 0.94±0.12 | 0.27±0.05 | 28.44±8.67 |
| 17 | 211 | 74.58±8.29 | 3.97±0.71 | 0.49±0.14 | 47.64±6.78 |
| 18 | 212 | 53.59±4.39 | 0.99±0.05 | 16.43±2.76 | 45.45±11.36 |
| 19 | 214 | 49.43±2.37 | 0.82±0.09 | 38.84±9.64 | 80.19±8.32 |

We have been successful in improving the affinity of the designed MTDLs towards NMDAR without losing their potency towards AChE (Fig. 2A and 2D, Table 1). Indeed AChE inhibitory activity was improved for some of the MTDLs (Compounds 201, 203, 205, 208, 209, 212 and 214). Although the MTDLs 203, 205, 208 and 211 exhibited slight toxicity in HepG2 cells (Fig. 2G), they are better than tacrine. These MTDLs also exhibited more potency towards AChE and NMDAR when compared to tacrine. Especially for MTDLs 201 and 208, the NMDAR and AChE inhibitory activities were improved when compared to tacrine. Hence, we assume that these compounds would be effective in inhibiting their targets at lower doses than tacrine and thereby may have lesser hepatotoxicity under *in vivo* conditions.

Our MTDLs reported in the current study are more potent than other FDA approved AChEI such as Rivastigmine (*IC*_*50*_=4.15 μM) [13] and Galantamine (*IC*_*50*_=15.06 μM) [82]. Hence, we believe that at reduced doses of these MTDLs, they could still be therapeutically effective. Moreover, increase in the efficacy of NMDAR inhibition could contribute to the multi targeting effect to achieve overall therapeutic outcome. Inhibition of BACE-1 activity is an additional desirable property for the MTDLs. Hence, we expect that the MTDLs reported in the current study will be beneficial for the treatment of AD as they can act on multiple pathways that underlie cognitive decline and neuronal loss towards achieving disease modification.

Although the etiology varies among different neurodegenerative diseases, neuronal injury and subsequent neuronal death is common. This is primarily caused by excessive activation of NMDAR and hence NMDAR antagonists might be useful in various neurodegenerative conditions. We also expect that these MTDLs would be beneficial in acute conditions such as ischemia [28], stroke [29] and TBI [30] in which NMDAR hyperactivity causes significant damage. In acute conditions, since the NMDAR antagonists are used for relatively shorter duration, the risk of side effects would be lesser and hence higher doses might be permissible. Our biochemical and cytological investigations revealed that glutamate induced excitotoxicity and the consequent death of neurons in culture were prevented by MTDLs such as 14, 201, 208, 210 and 211. Also, these compounds were neuroprotective (Fig. 3B) at the concentration at which they showed effective inhibition of heterologously expressed NMDAR (Fig. 2D) indicating that their mechanism of neuroprotection might be driven by NMDAR inhibition.

Additionally, the *in vivo* neuroprotection studies of 201 and 208 indicated that they can effectively rescue brain function from MSG mediated impairment. Although 201 is a moderate inhibitor of NMDAR compared to 208, it was efficient in neuroprotection both *in vivo* and *in vitro.* Previous studies demonstrated the neuroprotective properties of aryl azoles and have shown that methyl pyrazole moiety can favor neuroprotective property[83]. Hence, we presume that methyl-pyrazole moiety also might have contributed to the neuroprotective property of 201 apart from NMDAR inhibition. The neuroprotective action of 208 might be solely related to NMDAR inhibition since it a potent NMDAR antagonist.

MSG has been shown to induce a series of behavioral impairments and brain lesions in various experiments by overactivating glutamate receptors [75–79]. MSG treatment protocol that we used created a model of excitotoxic stress of a subacute and chronic nature. We believe the protection offered by 201 and 208 in the MSG treated rats could be the result of the NMDAR inhibitory activities of these compounds. While the *in vivo* efficacy of these compounds needs to be tested in more animal models of cholinergic and/or glutamatergic deficiencies, our data show that the newly designed MTDLs are indeed acting in the animal model as predicted by the modeling and *in vitro* studies.

## Conclusion

Due to the multifactorial etiology of neuropsychiatric diseases, MTDLs have been suggested as better disease modifying agents than single target directed drugs. It is widely accepted that these MTDLs not only can earnestly ameliorate the symptoms but also modify the disease. Tacrine, the first approved drug for AD, had hepatotoxicity. In spite of this, tacrine moiety has served as a template for the designing of hybrids/MTDLs. Though tacrine has been reported as a potent AChE inhibitor and weak NMDAR antagonist, its binding mode was unknown. Hence in this study, we demonstrated the binding modes of tacrine towards AChE and NMDAR through molecular modeling. With an aim to design potential MTDLs against AChE and NMDAR, we rationally designed 75 tacrine derivatives based on the predicted binding mode of tacrine. Among them, 19 molecules were chemically synthesized and were evaluated *in vitro* against both of these targets. We found that the derivatization improved AChE inhibitory potential for few compounds like, 13, 17, 201, 203, 205, 208, 209, 211, 212 and 214. The derivatization also improved the antagonistic potential towards NMDAR for compounds like 10, 14, 107, 201, 203, 204, 205, 206, 208, 209, 210, 211, 212 and 214. Based on the inhibition data, we tried to generate an SAR for the designed compounds. We found that substitutions at R1 and R4 positions together, reduce NMDAR and AChE inhibitory activity. The aromatic/hetero aromatic substitutions at R4 position of tacrine are key determinants to improve the antagonistic potential towards NMDAR. These SAR would be useful for future discovery of potent tacrine derived MTDLs. As an additional advantage, we observed BACE-1 inhibitory activity for compounds 10, 204, 206, 208 and 210. The preliminary assessment of hepatotoxicity of the MTDLs on HepG2 cell line suggested that derivatization lowered the toxic nature of the compounds when compared to the parent compound, tacrine. A few MTDLs were tested for their effect on rat primary cortical neurons subjected to glutamate induced excitotoxicity and were found to be neuro-protective. Further, *in vivo* studies of the selected tacrine derived MTDLs 201 and 208 using a rat model of MSG induced excitotoxicity showed that these compounds were able to protect the animals against MSG induced behavioral impairment thus demonstrating their efficacy *in vivo*. Our *in silico* ADME predictions of 201 and 208 suggested that they are drug like molecules with promising therapeutic and acceptable pharmacokinetic properties encouraging further *in vivo* studies. Based on the entire study, we suggest that tacrine derived MTDL, 201 is a potential drug candidate for further evaluation *in vivo* while the other derivatives need more *in vitro* and *in vivo* investigations.

## Materials and Methods

### Computational approaches

#### Modeling of NMDAR, structure-based design of tacrine derivatives and their ensemble docking towards NMDAR and AChE

Extensive molecular modeling studies were performed to model human NMDAR (h-NMDAR), to understand the binding mode of tacrine towards AChE and NMDAR, to design tacrine derived MTDLs and to predict the binding affinity of the designed inhibitors against NMDAR and AChE.

#### Coarse-grained modeling of NMDAR

Crystal structure of GluN1/GluN2B delta-ATD NMDAR from *Xenopus laevis* in complex with MK-801 (NMDAR antagonist) (PDB ID: 5UN1) [84] was used for our modeling studies. Since, MK-801 is an NMDAR antagonist, its binding site in the pore was taken as the spatial reference for the docking of tacrine. Initially, the missing regions in GluN1/GluN2B were patched and mutations were replaced with original human NMDAR amino acids using structure prediction wizard of Schrödinger. We employed an extensive loop refinement to optimize the newly constructed regions. A systematic coarse-grained modeling using normal mode analysis (NMA) and elastic network modeling (ENM) was performed on the modeled structure to address the structural dynamics of NMDAR. Programs such as Phenix SCEDS, Bio3D, Schrödinger and web servers such as WEBnma and iMod were used for the NMA and elNémo was used for ENM [85–89]. Majority of the programs except Phenix SCEDS provided multiple output structures in the form of poly alanine C-alpha traces. Amino acid backbone structures were built from the C-alpha traces by optimizing the main chain hydrogen-bonding networks using REMO algorithm [90]. Finally, polyalanine-independent conformational conversion of these structures was performed using the ‘Mutate residue range’ option using Coot. All output structures were clustered based on the global RMSD and one candidate structure was chosen from each of the clusters and was prepared for ensemble docking.

#### Preparations of tacrine and tacrine derived MTDLs for docking studies

Compounds such as tacrine and its derivatives were prepared using LigPrep module, Schrödinger and was used for molecular docking. LigPrep produced energy-minimized and protonated structures at pH 7.0 ±2 with different ionization states, tautomerism, stereo isomerism and ring conformations. Optimized potential for liquid simulations (OPLS3) force field was used for energy minimization of the ligand structure [91].

#### Ensemble docking of tacrine with NMDAR

Prior to ensemble docking, the candidate receptor structures were prepared using protein preparation wizard of Schrödinger. During the preparation, hydrogen atoms were added to the polar groups, bond orders were corrected and a short energy minimization was performed with an RMSD cutoff of 0.30 Å using OPLS-3 force field. By specifying the spatial reference of MK-801, a grid of size 20 Å was prepared on the four ensemble structures. The size of the grid was designed in such a way that it will cover the whole channel and hence all possible binding sites in the channel. Finally, the prepared ligands were docked in the grid using standard precision and induced fit docking method.

#### Ensemble docking of tacrine with AChE

In order to understand how multiple conformations of AChE affect the binding of tacrine and its derivatives, ensemble docking was carried out. Four different human AChE (hAChE) structures were taken for docking studies. These structures differ in the conformation of an active site residue, Y337. Out of four structures, one was the apo structure (PDB ID: 4EY4) [62] and the others were in complex with donepezil (PDB ID:4EY7) [62], huperzine (PDB ID:4EY5) [62] and 9-aminoacridine (PDB ID:6O4X) [92]. These hAChE structures were prepared for docking as described above after removing the crystallographic water molecules.

#### Structure based designing of tacrine derivatives, ensemble docking towards NMDAR and AChE and their binding energy calculations

In order to design MTDLs, the lowest energy receptor bound conformation of tacrine was used. Previous studies which explained the binding modes of different tacrine derivatives against AChE [48] also served as a potential references for designing MTDLs. We generated several tacrine derived MTDLs using a rational design approach by taking into consideration the conformations of residues surrounding the bound tacrine. Standard precision and induced fit docking method was used to predict the binding mode of the designed ligands towards both targets. After docking, the binding energy was estimated for each ligand complex using prime MM-GBSA method and structure activity relationships were deduced computationally.

#### Prediction of ADME properties

For computational assessment of absorption, distribution, metabolism and excretion (ADME) properties of the compounds, we calculated physico-chemical parameters of the selected MTDLs using QikProp, Schrödinger and SwissADME [93]. Parameters important for CNS activity and blood-brain barrier (BBB) permeability were mainly assessed for selected molecules using QikProp, SwissADME and online BBB predictor [94].

#### Chemical synthesis

In order to validate our *in silico* findings, promising MTDLs were chemically synthesized. All experiments were set-up on fume hoods and were carried out under nitrogen atmosphere in Schlenk tubes, unless otherwise noted. All solvents and reagents were procured from commercially available sources like Sigma-Aldrich, Combi-bolcks and Spectrochem. Commercially available pre-packed silica gel (230-400 mesh) plugs were used for column chromatography. All of the synthesized molecules were purified using solvents such as hexane, ethyl acetate, dichloromethane and methanol. Isolated yields correspond to products of >95% purity for the synthesized compoundsas determined by LC-MS and NMR.^1^H NMR was recorded on Bruker 400MHz AVANCE series or Bruker300 MHz DPX Spectrometer with CD_3_OD or DMSO-D_6_ as the solvent. All NMR chemical shifts were reported in parts per million (ppm), all coupling constants were reported in Hertz (Hz) and tetramethylsilane was used as internal standard for ^1^H NMR. Multiplicities are abbreviated as follows: singlet (s), doublet (d), triplet (t), quartet (q), doublet-doublet (dd), multiplet (m), and broad (br). Liquid chromatography-mass spectrometry (LC-MS) was used for reaction monitoring and determination of product mass on Agilent 1100 Series LC/MSD mass spectrometer. The compound numbers shown in parenthesis (Table S1) indicate the compound numbering according to our patent application (IPO-201841015699).

#### General procedure for the synthesis of 6-bromo-1,2,3,4-tetrahydroacridin-9-amine (Compound 3)

To a solution of 2-amino-4-cyano-benzonitrile (1) (5.0 g, 0.0253 mol, 1.0 equiv) in anhydrous toluene was added boron trifluoride etherate (3.76 mL, 0.0304 mol, 1.2 equiv) slowly at room temperature. The reaction mixture was cooled to 0°C followed by the addition of cyclohexanone (2) (3.9 mL, 0.0379, 1.5 equiv) and the reaction mixture was heated to 100°C for 4 h. After completion of reaction as monitored by TLC, the reaction mixture was quenched with aq.NaOH solution upto pH=10 and the reaction mixture was diluted with ethyl acetate. The organic layers were separated, dried over sodium sulphate and concentrated. The crude product was recrystallized from dichloromethane to obtain 6-bromo-1,2,3,4-tetrahydroacridin-9-amine (3) (5.8 g, 82%) as pale-yellow solid. ^1^H NMR: DMSO-*d6* (400 MHz): δ 8.13 (d, 1H, *J*= 8.8 Hz), 7.81 (s, 1H), 7.40-7.37 (m, 1H), 6.50 (bs, 2H), 2.82-2.79 (m, 2H), 2.55-2.52 (m, 2H), 1.82-1.80 (m, 4H).LCMS: m/z 279.0 (M+2), Analysis calculated for C_13_H_13_BrN_2_

#### General procedure for the synthesis of 9-amino-6-(1-methyl-1H-pyrazol-4-yl)-1,2,3,4-tetrahydroacridinium (Compound 201)

To a solution of 6-bromo-1,2,3,4-tetrahydroacridin-9-amine (3) (100 mg, 0.36 mmol, 1.0 equiv) in 1,4-dioxane (4 mL) and water (0.5 mL) were added 1-methyl pyrazole 4-boronic ester (4a) (90 mg, 0.434 mmol, 1.2 equiv) and sodium carbonate (57 mg, 0.54 mmol, 1.5 equiv). The reaction mixture was degassed for 10 min and Pd(PPh_3_)_4_ (41.0 mg, 0.036 mmol, 0.1 equiv) was added. The reaction mixture was heated to 110°C for 2h in a sealed tube. The reaction mixture was filtered through celite, washed with ethyl acetate (50 ml). The filtrate was diluted with water (50 mL) and extracted by ethyl acetate (3 × 50 mL). The organic layers separated were dried over anhydrous sodium sulphate and were concentrated under vacuum. The crude product was purified by column chromatography using 10% methanol in dichloromethane as eluent to give 6-(1-methyl-1H-pyrazol-4-yl)-1,2,3,4-tetrahydroacridin-9-amine (Compound 201) (50 mg, 50% yield) as off white solid.^1^H NMR: NMR: MeOD (400 MHz): δ 8.13-8.10 (m, 2H), 7.97 (s, 1H), 7.81 (s, 1H), 7.66 (dd, 2H, J = 8.4, 1.6 Hz), 3.98 (s, 3H), 2.97-2.94 (m, 2H), 2.65-2.62 (m, 2H), 1.97-1.95 (m, 4H). LCMS: m/z 279.1 (M+1), Analysis calculated for C_17_H_18_N_4_.

#### 6-(pyrimidin-5-yl)-1,2,3,4-tetrahydroacridin-9-amine (Compound 203)

Compound 203 was synthesized in a similar way as 201 but using pyrimidin-5-ylboronic acid (4b) (54 mg, 0.434 mmol, 1.2 equiv) instead of 4a; Yield: 55 mg, 55%; ^1^H NMR: MeOD (400 MHz): δ 9.18 (s, 3H), 8.22-8.20 (m, 1H), 8.00 (m, 1H), 7.69-7.66 (dd, 1H, *J*=8.8, 1.8 Hz), 2.97-2.94 (m, 2H), 2.66-2.63 (m, 2H), 1.96-1.93 (m, 4H). LCMS: m/z 277.2 (M+1), Analysis calculated for C_17_H_16_N_4_.

#### 6-(1H-pyrazol-3-yl)-1,2,3,4-tetrahydroacridin-9-amine (Compound 204)

Compound 204 was synthesized in a similar way as 201 but using (1H-pyrazol-3-yl) boronic acid (4c) (49 mg, 0.434 mmol, 1.2 equiv) instead of 4a; Yield: 48 mg, 50%; ^1^H NMR: MeOD (400 MHz): δ 8.10-8.05 (m, 2H), 7.82-7.79 (m, 1H), 7.71 (m, 2H), 6.81(d, 1H, *J*=2.0 Hz), 2.96-2.93 (m, 2H), 2.66-2.63 (m, 2H), 1.95-1.90 (m, 4H). LCMS: m/z 265.2 (M+1), Analysis calculated for C_16_H_16_N_4_.

#### 4-(9-amino-5,6,7,8-tetrahydroacridin-3-yl)benzonitrile (Compound 205)

Compound 205 was synthesized in a similar way as 201 but using (4-cyanophenyl) boronic acid (4d) (64 mg, 0.434 mmol, 1.2 equiv) instead of 4a; Yield: 60 mg, 55%; ^1^H NMR: DMSO (300 MHz): δ 8.29-8.26 (m, 2H), 8.19-8.16 (m, 1H), 8.00-7.99 (m, 1H), 7.86-7.83 (m, 1H), 7.72-7.67 (m, 2H), 6.44 (bs, 2H), 2.90-2.80 (m, 2H), 2.72-2.68 (m, 2H), 1.90-1.87 (m, 4H). LCMS: m/z 300.2 (M+1), Analysis calculated for C_20_H_17_N_3_.

#### 6-(4-(trifluoromethoxy)phenyl)-1,2,3,4-tetrahydroacridin-9-amine (Compound 206)

Compound 206 was synthesized in a similar way as 201 but using (4-(trifluoromethoxy)phenyl) boronic acid (4e) (90 mg, 0.434 mmol, 1.2 equiv) instead of 4a; Yield: 59 mg, 45%; ^1^H NMR: MeOD (400 MHz): δ 8.17 (d, 1H, *J*= 8.8 Hz), 7.96-7.94 (m, 1H), 7.86-7.84 (dd, 2H, *J*=6.4, 2.0 Hz), 7.69-7.66 (dd, 1H, *J*=4.8, 2.0 Hz), 7.43-7.41 (m, 2H), 2.98-2.95 (m, 2H), 2.67-2.64 (m, 2H), 2.01-1.90 (m, 4H). LCMS: m/z 359.0 (M+1), Analysis calculated for C_20_H_17_F_3_N_2_O.

#### 6-(2-fluorophenyl)-1,2,3,4-tetrahydroacridin-9-amine (Compound 208)

Compound 208 was synthesized in a similar way as 201 but using (2-fluorophenyl) boronic acid (4g) (61 mg, 0.434 mmol, 1.2 equiv) instead of 4a; Yield: 63 mg, 60%; ^1^H NMR: MeOD (400 MHz): δ 8.42 (d, 1H, *J*= 8.8 Hz), 7.94 (s, 1H), 7.94-7.80 (m, 1H), 7.67-7.72 (m, 1H), 7.60-.7.49 (m, 1H), 7.40-7.37 (m, 1H), 7.37-7.25 (m, 1H), 3.05-3.02 (m, 2H), 2.68-2.65 (m, 2H), 2.03-1.98 (m, 4H). LCMS: m/z 293.2 (M+1), Analysis calculated for C_19_H_17_FN_2_.

#### 6-(4-fluorophenyl)-1,2,3,4-tetrahydroacridin-9-amine (Compound 209)

Compound 209 was synthesized in a similar way as 201 but using 2-(4-fluorophenyl)=4,4,5,5-tetramethyl 1,3,-dioxaborolane (4h) (96 mg, 0.434 mmol, 1.2 equiv) instead of 4a; Yield 72 mg, 68%; ^1^H NMR: MeOD (400 MHz): δ 8.37 (d, 1H, *J*= 8.4 Hz), 7.87-7.81 (m, 2H), 7.79-7.77 (m, 2H), 7.70-7.61 (m, 1H), 7.58-7.55 (m, 1H), 7.29-7.24 (m, 2H), 3.03-3.02 (m, 2H), 2.63-2.61 (m, 2H), 2.05-1.90 (m, 4H). LCMS: m/z 293.2 (M+1), Analysis calculated for C_19_H_17_FN_2_.

#### 6-(4-(methylthio)phenyl)-1,2,3,4-tetrahydroacridin-9-amine (Compound 210)

Compound 210 was synthesized in a similar way as 201 but using 4,4,5,5-tetramethyl-2-(4-(methylthio) phenyl)-1,3,2-dioxaborolane (4i) (109 mg, 0.434 mmol, 12 equiv) instead of 4a; Yield: 64 mg, 55%;^1^H NMR: MeOD (400 MHz): δ 8.39 (d, 1H, *J*= 8.0 Hz), 7.91 (m, 2H), 7.43 (d, 2H, J=8.8), 7.42 (d, 2H, J=8.8), 3.05-3.02 (m, 2H), 2.67-2.64 (m, 2H), 2.56 (s, 3H), 2.01-2.00 (m, 4H). LCMS: m/z 312.2 (M+1), Analysis calculated for C_20_H_20_N_2_S.

#### 6-(4-(trifluoromethyl)phenyl)-1,2,3,4-tetrahydroacridin-9-amine (Compound 211)

Compound 211 was synthesized in a similar way as 201 but using (4-(trifluoromethyl)phenyl) boronic acid (4j) (83 mg, 0.434 mmol, 1.2 equiv) instead of 4a; Yield: 75 mg, 61%; ^1^H NMR: MeOD (400 MHz) δ 8.47 (d, 1H, *J*= 8.8 Hz), 8.01-7.92 (m, 4H), 7.88-7.86 (m, 2H), 3.10-3.06 (m, 2H), 2.72-2.65 (m, 2H), 2.10-1.99 (m, 4H). LCMS: m/z 343.0 (M+1), Analysis calculated for C_20_H_17_F_3_N_2_.

#### 6-(furan-3-yl)-1,2,3,4-tetrahydroacridin-9-amine (Compound 212)

Compound 212 was synthesized in a similar way as 201 but using 2-(furan-3-yl)-4,4,5,5-tetramethyl-1,3,2-dioxaborolane (4k) (84 mg, 0.434 mmol, 1.2 equiv) instead of 4a; Yield: 35 mg, 37%; ^1^H NMR: MeOD (400 MHz): δ 8.33 (d, 1H, *J*=8.8 Hz), 8.21 (s, 1H), 7.87-7.81 (m, 2H), 7.65-7.55 (m, 2H), 3.04-3.01 (m, 2H), 2.66-2.63 (m, 2H), 2.03-1.99 (m, 4H). LCMS: m/z 265.2 (M+1), Analysis calculated for C_17_H_16_N_2_O.

#### 6-(pyridin-3-yl)-1,2,3,4-tetrahydroacridin-9-amine (Compound 214)

Compound 214 was synthesized in a similar way as 201 but using 3-(4,4,5,5-tetramethyl-1,3,2-dioxaborolan-2-yl)pyridine (4m) (89 mg, 0.434 mmol, 1.2 equiv) instead of 4a; ^1^H NMR: MeOD (400 Hz): 9.44 (s, 1H), 9.14 (d, 1H, *J*= 8.0 Hz), 9.01 (d, 1H, *J*= 6.4 Hz), 8.62 (d, 1H, *J*= 8.8 Hz), 8.31 (m, 1H), 8.24 (s, 1H), 8.09 (d, 1H, *J*= 8.8 Hz), 3.11-3.09 (m, 2H), 2.70-2.67 (m, 2H), 2.03-2.02 (m, 4H); LCMS: m/z 276.0 (M+1), Analysis calculated for C_18_H_17_N_3_.

#### General procedure for the synthesis of Ethyl 9-amino-1,2,3,4-tetrahydroacridine-2-carboxylate (Compound 16)

To a solution of 2-amino benzonitrile (1a) (5.0 g, 0.0253 mol, 1.0 equiv) in anhydrous toluene was added boron trifluoride etherate (3.76 mL, 0.0304 mol, 1.2 equiv) slowly at room temperature. The reaction mixture was cooled to 0°C followed by the addition of ethyl-4-oxocyclohexanecarboxylate (2a) (6.4 mL, 0.0379mol, 1.5 equiv) and the reaction mixture was heated to 100°C for 4 h. After completion of reaction as monitored by TLC, the reaction mixture was quenched with aqueous sodium hydroxide solution at pH=10 and the reaction mixture was diluted with ethyl acetate. The organic layers were separated, dried over sodium sulphate and concentrated. The crude product was recrystallized from dichloromethane to obtain ethyl 9-amino-1,2,3,4-tetrahydroacridine-2-carboxylate (Compound 16) (7.0 g, 85%) as off white solid [48]. ^1^H NMR (300 MHz, DMSO-*d6*): δ 8.14 (d, 1H,*J* = 8.37 Hz), 7.61 (d, 1H, *J* = 8.31 Hz), 7.48 (t, 1H, *J* = 6.9 Hz), 7.27 (t, 1H, *J* = 6.93 Hz), 6.43 (s, 2H), 4.18-4.08 (m, 2H), 2.87-2.82 (m, 4H), 2.72-2.62 (m, 1H), 2.26e2.12 (m, 1H), 1.88e1.82 (m, 1H), 1.23 (t, 3H,*J* = 7.11 Hz). LCMS: m/z calculated for C_16_H_18_N_2_O_2_: 270.33; Observed mass: 271.2 (M + H).

#### Ethyl 9-amino-6-bromo-1,2,3,4-tetrahydroacridine-2-carboxylate (Compound 5)

Compound 5 was synthesized in a similar way as 16 but using 4-bromo-2-amino benzonitrile. ^1^H NMR (400 MHz, DMSO-*d6*): δ 8.13 (d, 1H, *J* = 8.96 Hz), 7.79 (s, 1H), 7.40 (dd, 1H, *J* = 8.92, 8.92 Hz), 6.61 (s, 2H), 4.19-4.07 (m, 2H), 2.94-2.85 (m, 4H), 2.68-2.61 (m, 1H), 2.15-2.11 (m, 1H), 1.89-1.80 (m, 1H), 1.24 (t, 3H,*J* = 7.12 Hz). LCMS: 351.7 (M+2), m/z calculated for C_16_H_17_BrN_2_O_2_.

#### General procedure for the synthesis of Ethyl-9-amino-6-(1-methyl-1H-pyrazol-4-yl)-1,2,3,4-tetrahydroacridine-2-carboxylate (Compound 10)

To a solution of ethyl 9-amino-6-bromo-1,2,3,4-tetrahydroacridine-2-carboxylate **(5)** (1.0 g, 2.87 mmol) in 1,4-dioxane (10 mL) and water (1 mL), 1-methyl pyrazole 4-boronic ester (4a) (0.9 g, 4.31 mmol) and sodium carbonate (0.6 g, 5.74 mmol) were added. The reaction mixture was degassed for 10 min and Pd(PPh_3_)_4_ (0.33 g, 0.287 mmol) was added. The reaction mixture was heated to 110°C for 2h in a sealed tube. The reaction mixture was filtered through celite, the filtrate was diluted with water (200 mL) and was extracted with ethyl acetate (3×200 mL). The organic layers separated were dried over anhydrous sodium sulphate and were concentrated under vacuum. The residue obtained was recrystallized in dichloromethane to afford compound 10 (0.52 g, 52%) as white solid [48]. ^1^H NMR (400 MHz, DMSO-*d6*): δ 8.26 (s, 1H), 8.16 (d, *J* = 8.0 Hz, 1H), 7.99 (m, 1H), 7.82 (m, 1H),7.55 (m, 1H), 6.57 (bs, 2H), 4.15 (m, 2H), 3.89 (s, 3H), 2.88 (m, 4H), 2.69 (m, 1H), 2.16 (m, 1H), 1.90 (m, 1H), 1.26 (t, *J* = 7.12, 3H). LCMS:336.2 (M+H) m/z calculated for C_19_H_21_N_5_O.

#### Synthesis of ethyl 9-amino-6-(4-fluorophenyl)-1,2,3,4-tetrahydroacridine-2-carboxylate (Compound 107)

Compound 107 was synthesized in a similar way as 10 but using 2-(4-fluorophenyl)-4,4,5,5-tetramethyl-1,3,2 dioxaborolane (4h); ^1^H NMR (400 MHz, MeOD): δ 8.14 (d, J=8.8 Hz, 1H), 7.91 (m, 1H), 7.80-7.75 (m, 2H), 7.58 (dd, J=8.0, 2.0 Hz, 1H), 7.24 (m, 2H), 4.22 (m, 2H), 3.05-3.01 (m, 2H), 2.99-2.90 (m, 2H), 2.89-2.82 (m, 1H), 2.33-2.29 (m, 1H), 2.6-1.98 (m, 1H), 1.32 (t, J=7.2 Hz, 3H); LCMS: 365.2 (M+H) m/z calculated for C_22_H_21_FN_2_O_2_.

#### General procedure for the synthesis of 9-Amino-N-methyl-6-bromo-1,2,3,4-tetrahydroacridine-2-carboxamide (Compound 13)

To a solution of compound 5 (100 mg, 0.37 mmol) in methanol (4 ml) aq. ammonia (1 ml) was added and the reaction mixture was heated to 60°C for 2h. The reaction mixture was concentrated to dryness and was recrystallized from dichloromethane to give compound 13 [48]. White solid; Yield 80 mg, 80%; ^1^H NMR (400 MHz, DMSO-d6): **δ** 8.15-8.12 (m, 1H), 7.94 (d, J = 4.4 Hz, 1H), 7.81 (m, 1H), 7.42 (d, J = 8.8 Hz, 1H), 6.68 (bs, 3H), 2.88 (s, 3H), 2.76-2.68 (m, 3H), 2.15-2.12 (m, 1H), 2.01-1.98 (m, 1H), 1.91-1.81 (m, 2H). LCMS: m/z calculated for C_15_H_17_N_3_O: 225.14; Observed mass: 226.2 (M + H).

#### 9-amino-N-methyl-6-(1-methyl-1H-pyrazol-4-yl)-1,2,3,4-tetrahydroacridine-2-carboxamide (Compound 14)

Compound 14 was synthesized in a similar way as 13 but using compound 10; Yield: 50 mg, 92 % as a white solid.^1^H NMR (400 MHz, DMSO-d6): δ 8.37 (s, 1H), 8.16 (d, J = 8.4 Hz, 1H), 7.99 (s, 1H), 7.95 (m, 1H), 7.81 (s, 1H), 7.56 (d, J = 8.0 Hz, 1H), 6.67 (bs, 2H), 3.89 (s, 3H), 2.89 (m, 2H), 2.60 (m, 4H), 2.33 (m, 1H), 2.15 (m, 1H), 1.91 (m, 1H), 1.81 (s, 1H). LCMS: 351.4 (M+H) m/z calculated for C_20_H_22_N_4_O_2_.

#### General procedure for the synthesis of 9-amino-1,2,3,4-tetrahydroacridine-2-carbohydrazide (Compound 17)

To a solution of compound 16 (100 mg, 0.37 mmol) in methanol (4 ml), hydrazine hydrate (1 ml) was added and the reaction mixture was refluxed for 3h at 70°C. The reaction mixture was concentrated and was recrystallized from dichloromethane to give compound 17 [48]. Brown solid; Yield 80 mg, 85%;^1^H NMR (400 MHz, MeOD): δ 8.28-8.20 (m, 1H), 7.82-7.80 (m, 1H), 7.74-7.71 (m, 1H),7.69-7.58 (m, 1H), 3.09-3.03 (m, 2H), 2.91-2.80 (m, 3H), 2.26-2.22 (m, 1H), 2.06-2.03 (m, 1H); LCMS: m/z calculated for C_14_H_16_N_4_O: 256.13; Observed mass: 257.2 (M + H);

#### 9-Amino-6-bromo-1,2,3,4-tetrahydroacridine-2-carbohydrazide (Compound 8)

Compound 8 was synthesized in a similar way as 17 but using compound 5; White solid; Yield 70 mg, 73%; ^1^H NMR (400 MHz, DMSO-d6): δ 8.15 (s, 1H), 8.11(d, J = 8.8 Hz, 1H), 7.79 (d, J = 2.0 Hz, 1H), 7.40-7.37 (m, 1H), 6.56 (s, 2H), 4.24 (bs, 2H), 2.91-2.77 (m, 2H), 2.72-2.67 (m, 2H), 1.96 (m,1H), 1.83 (m, 1H); LCMS: m/z calculated for C_14_H_15_BrN_4_O: 334.04; Observed mass: 336.2 (M+2);

### *In vitro* studies

To understand the pharmacological properties of the synthesized MTDLs, we performed *in vitro* activity assays of cholinesterases, NMDAR and BACE-1.

#### *In vitro* cholinesterase (AChE and BChE) inhibition assays

Enzymatic assays for AChE were carried out in a 96-well plate using AMPLITE™ AChE assay kit (AAT Bioquest, Inc., Sunnyvale, CA, USA) which consists of AChE from *Electrophorus electricus* (electric eel) (structural similarity towards human AChE is ≥ 90 %) [95, 96], assay buffer (pH 7.4), 5,5-dithiobis-(2-nitrobenzoic acid) (DTNB, known as Ellman’s reagent) and the substrate acetylthiocholine (AChT). The assay system works on the basis of Ellman’s method [97]. Thiocholine, produced by the action of AChE on acetylthiocholine, forms a yellow colored product with DTNB. The quantity of the colored product measured at 410 ± 5 nm, is proportional to enzyme activity. The native enzyme reaction was carried out by mixing enzyme (0.3U) with 125 μM AChT (in ddH_2_O) and 125 μM DTNB (in assay buffer). The same experiment was repeated in the presence of the selected MTDLs. The MTDLs, were dissolved in 0.01 % DMSO and were pre-incubated with AChE at room temperature for 20 minutes prior to the addition of reaction mixture containing AChT and DTNB. The OD at 405 nm of the reaction mixture in the presence and absence of compounds was plotted against time and from there the relative activity of enzyme in the presence of compounds was calculated. The half maximal inhibitory concentration (*IC*_*50*_) was determined for the MTDLs by performing the assay at different inhibitor concentrations.

The inhibitory effect of all the compounds against butyrylcholinesterase (BChE) was also determined in a similar way. In this experiment, BChE from equine serum (0.3U), butyrylthiocholine (500 μM) and DTNB (500 μM) in PBS (pH 7.4) were used for the inhibition assays. In all the experiments, the *IC*_*50*_ values are presented as the mean ± standard deviation of at least three separate experiments.

#### Effect of MTDLs on NMDA receptor activity

To check the effect of MTDLs on NMDAR activity a cell-based assay system that works on the basis of protein-protein interaction between NMDAR and α-CaMKII was used [66]. The plasmids coding for NMDAR subunits, GluN1 and GluN2B, and α-CaMKII tagged with GFP (GFP-α-CaMKII) were co-transfected into HEK-293 cells and the activity of GluN2B containing NMDAR was detected. In order to perform the assay, following procedures were performed.

#### Preparation and transfection of plasmids coding for GluN1, GluN2B and GFP-α-CaMKII to HEK-293 cells

The plasmids for mammalian expression carrying the cDNAs for GluN1, GluN2B and GFP-α-CaMKII were prepared from bacterial cell using QIAGEN Midi Kit (Qiagen, USA). The plasmids were quantified using NanoDrop 2000 and their purity was checked on 1% agarose gel as reported previously [66]. These plasmids were further co-transfected into pre-grown HEK-293 cells using lipofectamine (Invitrogen) according to the manufacturer’s protocol. (Concentrations of the plasmids per well used for transfection: GluN1 and GluN2B - 0.35 μg each and GFP-α-CaMKII - 0.15 μg). HEK-293 cells were grown by seeding them on sterile 12 mm coverslips placed in 24-well plates (~1.5 × 10^4^ cells/well). About 18 hours after seeding, the cells were co-transfected with the plasmids. Upon terminating the transfection reaction after 5 hours with 20% fetal bovine serum (FBS), 20 μM MK-801was also added. The addition of MK-801 prevented cell death in transfected cells that can happen due to activation of NMDAR. The transfection solution was aspirated after 12 hours and 500 μL of fresh Dulbecco’s modified Eagle’s medium (DMEM) containing 10% FBS and 20 μM MK-801 was added. The cells were further incubated at 37°C for 24 hours.

#### NMDAR activity assay

About 24 hours after transfection, cells were washed twice, first with Hank’s balanced salt solution (HBSS) containing 1 mM HEPES and 0.5 mM EGTA (Solution Ι) and then with HBSS containing 1 mM HEPES (Solution ΙΙ). NMDAR was activated with Solution ΙΙΙ which contains NMDAR agonists such as glutamate (100 μM) and glycine (10 μM), along with Ca^2+^ (2 mM) in solution II. After 5 minutes of activation, cells were fixed with 4% paraformaldehyde (PFA) for 10-15 minutes and were washed thrice with PBS. The cover slip containing cells were mounted on a clean glass slide using glycerol:PBS (1:1) and were visualized using a fluorescence microscope. The activity of GluN2B containing NMDAR was detected based on translocation of GFP-α-CaMKII to GluN2B subunit in the endoplasmic reticulum (ER), nuclear membrane and plasma membrane. It was observed as redistribution of green fluorescence accumulation (punctae) at the membranes. To check the effect of MTDLs on NMDAR activity, the compounds at respective concentrations were included in both solution ΙΙ (pre-incubation for 5 min) and solution ΙΙΙ. In order to quantitate the NMDAR activity, the cells were counted using a fluorescence microscope (Leica DMI 4000B inverted microscope) at 40X magnification. The number of green fluorescent cells and green fluorescent cells with punctae were counted separately and the percentage of green fluorescent cells having a punctate pattern was calculated as [(number of punctate cells /total number of green fluorescent cells) × 100]. This number was taken as the efficiency of punctae formation or punctate cell count. For each slide, 5 or more fields were randomly selected and the average of punctate cell count for all the fields was estimated. Further, percent inhibition of activity in presence of the compounds with respect to the control was calculated. Final values were obtained from three independent experiments. For *IC*_*50*_ measurements, each estimation was done with 6 or more concentrations of the MTDLs. *IC*_*50*_ values from three such determinations were used to calculate mean±SD for each compound.

#### Effect of MTDLs on interaction between GFP-α-CaMKII and GluN2B

In order to study the effect of MTDLs on the interaction of GFP-α-CaMKII and GluN2B, HEK-293 cells stably expressing GFP-α-CaMKII and a construct having the GluN2B motif which binds to α-CaMKII, tagged with mitochondrial localization signal (MLS-NR2B) was used [66]. Since there are no functional NMDAR channels in the membrane, GFP-α-CaMKII was activated by Ca^2+^ influx through ionomycin, a Ca^2+^ ionophore. The activated GFP-α-CaMKII binds to MLS-NR2B and the interaction can result in the formation of perinuclear punctae of green fluorescence. To achieve this, cells grown on the coverslips were first washed with HBSS containing 1 mM HEPES and 0.5 mM EGTA for 5 minutes. Subsequently the cells were incubated with 250 μL HBSS containing 1 mM HEPES and 15 μM ionomycin with or without MTDL for 5 minutes. Subsequently, the cells were treated with HBSS containing 1 mM HEPES, 2 mM Ca^2+^ and 3 μM ionomycin with or without MTDL for 5 minutes. The cells were fixed using 4% PFA for 10 minutes and were washed twice with PBS for 5 minutes each. The coverslips were mounted onto glass slides and were used for imaging. Concentrations of the MTDLs which exhibited ≥ 70% inhibition on NMDAR activity were chosen for the studies.

#### Activity of selected MTDLs on BACE-1

BACE assay was carried out using a kit (PanVera, Invitrogen, USA) that utilizes fluorescence resonance energy transfer (FRET) based technology according to the manufacturer’s protocol. A peptide substrate based on amyloid precursor protein (Rh-EVNLDAEFK-Qu, where ‘Rh’ is Rhodamine and ‘Qu’ is quencher) was used. During enzymatic cleavage, due to the action of BACE-1, the fluorophore separates from the quencher and the substrate becomes highly fluorescent. Rate of increase in fluorescence is linearly proportional to enzyme activity. Assays were performed using 1U BACE-1 enzyme solution and 750 nM substrate in 50 mM sodium acetate buffer at pH 4.5 in a total volume of 100 μL. After incubating the reaction mixture for 60 minutes at 25 °C, under dark conditions, it was stopped using 2.5 M sodium acetate. Fluorescence was measured at 545 nm excitation and 585 nm emissions using a plate reader (Infinite^®^ 200 from TECAN). Approximately 50 μM of the selected MTDLs in 0.03% of DMSO were pre-incubated with enzyme solution for 15 minutes before the assay. Finally, % of residual BACE-1 activity was calculated in presence of compounds.

#### Cell viability and neuroprotection studies

To understand the toxicity profile of MTDLS, we have estimated their effect on the viability of HepG2 cells using MTT assay. The neuroprotective property of selected MTDLs against glutamate induced excitotoxicity was performed in primary rat cortical neuronal cells.

#### *In vitro* assay for hepatotoxicity of MTDLs

In the assay, the live cells are estimated by the reduction of MTT (pale yellow) into dark purple formazan crystals by the action of mitochondrial succinate dehydrogenase, which is monitored using spectrophotometer by measuring absorbance at 570 nm. Initially the HepG2 cells were maintained in DMEM supplemented with 10% FBS, 10,000 units/mL penicillin, 10 mg/mL streptomycin and 25 μg/mL amphotericin-B in T-25 flasks. Cells were trypsinized using 0.25% trypsin and were seeded onto 96-well plates (~15000 cells/well). After 24 hours, the medium was aspirated and was replaced with fresh medium containing varying concentrations of the selected MTDLs (10, 50, 100 and 300 μM). After growing the cells for 24 hours, 5 mg/mL MTT solution was added to the cells and was kept for an additional 2 hours. The formazan crystals formed were dissolved in 100 μL DMSO and the absorbance at 570 nm was measured using a multi-mode plate reader (Infinite^®^ 200 from TECAN). Cell viability is expressed as the percentage of viable cells compared to untreated cells.

### Experiments in primary neuronal cultures

#### Preparation of primary cortical neurons from E18 rat embryos

Primary cultures of cortical neurons were prepared from Wistar rat embryos at E18 stage. All animal studies were carried out at Rajiv Gandhi Centre for Biotechnology (RGCB), Thiruvananthapuram, according to the Institutional Animal Ethics Committee guidelines. The pregnant female rat was sacrificed, the uterus was carefully removed with the embryos and was placed in a 90 mm petri dish containing ice cold phosphate-buffered saline (PBS). Embryos were then removed from the uterus and the brain was dissected from each embryo. The cortical hemispheres were separated and the meninges were removed with the help of a stereo microscope. The tissues were minced gently in HBSS and were centrifuged at 345 g for 3 minutes. The tissue was then dissociated using 0.05% Trypsin-EDTA and DNase I (4000 U/mL) for 10 minutes. Action of trypsin on the tissue was arrested by addition of 10% FBS, followed by a wash with HBSS and centrifugation at 345 g for 3 minutes. Tissues were triturated to a single cell suspension using DMEM containing DNase I (4000 U/mL) and were washed twice with DMEM by centrifugation at 345 g for 5 minutes. Live cells were counted using trypan blue dye exclusion method and the cells were then seeded onto 24-well plates and 96-well plates, pre-coated with poly-D-lysine (100 μg/mL) and laminin (1 μg/mL), at a density of 1 × 10^5^ cells/well and 20 × 10^3^ cells/well respectively, in neurobasal medium (NBM) supplemented with 1X GlutaMAX, 1X antibiotic/antimycotic solution, 1X B27 supplement and ciprofloxacin (10 μg/mL). The plates were incubated at 37°C under 5% CO_2_. The cells were maintained in culture, by changing the media at regular intervals, till the day of experiment.

#### Induction of excitotoxicity in primary cortical neurons using glutamate

Glutamate-induced excitotoxicity was performed on primary cortical neurons that were maintained for nine days *in vitro* (DIV9) in culture. Prior to the experiment, the cells were washed with solution I (HBSS, 10 mM HEPES, pH 7.4 and 0.2 mM EGTA) for 10 minutes followed by treatment with solution II (HBSS, 10 mM HEPES, pH 7.4) for 10 minutes. Subsequently, these cells were incubated with solution III (HBSS, 10 mM HEPES, pH 7.4, 100 μM glutamate, 10 μM glycine, 1.2 mM CaCl_2_) for 60 minutes for induction of excitotoxicity. To check the activity of MTDLs, the compounds were included in solution II and III. MK-801 (20 μM) was used as a positive control in the experiment. After 60 minutes of glutamate treatment, solution III was replaced with NBM and the cells were maintained for 24 hours. The spent media was used for biochemical estimation of G6PD release and the cells were utilized for immunocytochemistry and DAPI staining by fixing them using 4% PFA. The cortical cells grown in 96-well plates were subjected to MTT assay after excitotoxicity treatment, which is explained in the section ‘MTT assay for measuring neuronal death’.

#### Glucose 6-phosphate dehydrogenase assay for measuring neuronal death

Since the damaged cells release G6PD to the surrounding medium, we quantified G6PD using Vybrant^TM^ cytotoxicity assay kit (Molecular Probes) and measured neuronal death. The assay kit utilizes the generated NADPH to reduce resazurin to red fluorescent resorufin (Abs/Em: 563/587 nm) by the action of diaphorase enzyme. The resulting fluorescence intensity is proportional to the amount of G6PD released into the medium which correlates with cell death. After excitotoxicity treatment, 50 μL of spent media was mixed with 50 μL of the reagent in a 96-well plate, incubated at 37^0^C for 40 minutes and fluorescence emission was measured at 587 nm. All necessary experimental controls and samples were assayed in duplicate.

#### MTT assay for measuring neuronal death

The cortical neurons subjected to excitotoxic treatments at DIV9 stage were subjected to MTT assay. Excitotoxicity was induced as explained in the section ‘Induction of excitotoxicity in primary cortical neurons using glutamate**’**except that glutamate treatment was given for 3 hours and the treatment solutions were replaced with NBM. The cells were maintained for another 24 hours and MTT assay has been carried out as explained in the section *‘In vitro* assay for hepatotoxicity of MTDLs’

#### Immuno and DAPI staining of primary cortical neurons

Immunostaining was carried out to visualize the distribution and localization of the neuronal marker protein, microtubule associated protein (MAP2) during excitotoxic conditions and DAPI was used as a nuclear counterstain. Cells after excitotoxicity treatment were fixed with PFA, washed thrice with PBS and were permeabilized and blocked simultaneously with PBS containing 0.2% triton X-100 and 3% BSA for 1 hour at room temperature. After removing the blocking solution, the cells were incubated with the primary antibody [MAP2 antibody −1:1000] diluted in PBS containing 2% BSA and 0.3% triton X-100 overnight at 4°C. The primary antibody was removed; the cells were washed thrice with PBS followed by incubation with secondary antibody conjugated to Cy3 at a dilution of 1:500, for 1-2 hours at room temperature. The cells were washed thrice with PBS before being stained with DAPI (1.8 μM) for 20 minutes. The cells were again washed thrice with PBS and the coverslips were mounted using fluoromount and were sealed with DPX. The slides were viewed using an epifluorescence microscope (Leica DMI 6000B inverted microscope) and the images were captured at 40X magnification. The percentage of viable cells and total number of cells were calculated from the DAPI stained nuclei in each field and percentage of MAP2 positive cells (calculation performed by counting MAP2 stained cells upon total number of DAPI stained cells in each field) were also calculated. All treatments were done in duplicates in each experiment. Values for 10 fields were averaged in each experiment. DMSO was included in the control.

#### Behavioral analysis using Morris water maze test

To understand the effect of MTDLs on *in vivo* excitotoxicity, we induced excitotoxic condition in rats by administering MSG. The treatment is expected to affect cognitive behavior of rats which was assessed using the MWM behavioral test. The performance in the test was compared between control rats and rats subjected to treatment of MTDLs. The experimental procedure is as follows:

Studies were conducted using 4-6 weeks old, male Wistar rats (~100g). MSG solutions were administered IP for 15 days. Every rat received two injections each day, an initial injection of the selected MTDL or vehicle followed after 30-45 min by saline or MSG (2-4 g/Kg body weight) in saline. The MWM test was started on the 11^th^ day of treatment, as described by Morris [98], with a slight modification. The water maze apparatus consists of a large circular pool (183 cm diameter, 64 cm height), with an escape platform (10 cm diameter, 35 cm height) and was filled with milky water to a level just above the platform. The rat was placed into the water, facing the tank wall (starting point) and was allowed to search for the platform for 60s (maximum trial time allowed) based on the visual cues present in the room. The trajectories of the rat were recorded using a video camera with EthoVision XT software (Noldus Information Technology). The time taken by each rat to reach the platform (escape latency) was measured using the software. If the rat finds the platform before 60s, it was allowed to stay on the platform for 5-10s and then was returned to its home cage. On the other hand, if the rat was unable to find the platform, it was physically placed on the platform for 10s and then was returned to its home cage. Each animal was given five trials per day (11^th^ to 15^th^ day of injections) with an interval of ~10-15s between each trial. For the initial screening of MTDLs, the rats were divided into the following groups, with three rats in each group (n=3): (i) Vehicle control (0.03% DMSO in saline), (ii) MSG (2 g/Kg body weight), (iii) compound 201 (3 mg/Kg body weight) control, (iv) compound 201 (3 mg/Kg body weight) and MSG, (v) compound 208 (1 mg/Kg body weight) control, (vi) compound 208 (1 mg/Kg body weight) and MSG, (vii) tacrine (5 mg/Kg body weight) control and (viii) tacrine (5 mg/Kg body weight) and MSG. For the subsequent confirmatory studies, the dose of MSG given was 4 g/Kg body weight and the n value was raised to at least 6.

#### Statistical analysis

The statistical analysis was done using one way ANOVA followed by Tukey’s post hoc test using Origin Pro 8 software. The significance level was set at p< 0.05. The quantitated values were represented as graphs with mean ± standard deviation (sd).

## Associated Content

Additional tables and figures are given in the supporting information.

## Abbreviations used

AD: Alzheimer’s disease
AChE: acetylcholinesterase
NMDAR: N-methyl-D-aspartate receptor
MTDLs: multi-target directed ligands
AChEIs: acetylcholinesterase inhibitors
TBI: traumatic brain injury
MWM: Morris water maze
RMSD: root mean square deviation
LBD: Ligand binding domain
TMD: transmembrane domain
ChE: Cholinesterases
Ca^2+^: calcium
BACE-1: β-site amyloid precursor protein cleaving enzyme
MAP2: microtubule associated protein 2
G6PD: glucose-6-phosphate dehydrogenase
IP: intraperitoneal
BBB: blood brain barrier
SVM: support vector machine
LiCABEDS: Ligand Classifier of Adaptively Boosting Ensemble Decision Stumps
SAR: structureactivity relationships
NMA: normal mode analysis
ENM: elastic network modeling
h-NMDAR: 
human NMDAR: hAChE, human AChE
ADME: absorption, distribution, metabolism and excretion
OPLS: Optimized potential for liquid simulations
ppm: parts per million
LC-MS: Liquid chromatography-mass spectrometry
DTNB: 5,5-dithiobis-(2-nitrobenzoic acid)
AChT: acetylthiocholine
BChE: Butyrylcholinesterase
DMEM: Dulbecco’s modified Eagle’s medium
HBSS: Hank’s balanced salt solution
PFA: paraformaldehyde
ER: endoplasmic reticulum
FRET: fluorescence resonance energy transfer
PBS: phosphate-buffered saline
NBM: neurobasal medium
SD: standard deviation

## Acknowledgements

C.R., K.V.D, M.K, L.K and R.S.J acknowledge the fellowships received from Indian Council of Medical Research (ICMR), Japan Society for the Promotion of Science (JSPS), Kerala State Council for Science Technology and Environment (KSCSTE), Council of Scientific and Industrial Research (CSIR) and Department of Science and Technology (DST) respectively. S.A greatfully acknowledges DST-SERB under young Scientist Scheme (SB/FT/CS-079/2014) for financial Support. Research funding received by R.V Omkumar from Rajiv Gandhi Centre for Biotechnology, Thiruvananthapuram is also acknowledged. We gratefully acknowledge the RIKEN ACCC for the supercomputing resources at the Hokusai Big Waterfall supercomputer facility and animal research facility of Rajiv Gandhi Centre for Biotechnology, Thiruvananthapuram for providing animals for conducting experiments. S.A. is thankful to VFSTR for analytical facilities at CoExAMMPC throughout the research work.

