## Supplemental data for "Neuroprotective derivatives of tacrine that target NMDA receptor and acetyl cholinesterase - Design, synthesis and biological evaluation"

### Supporting Information

| Title | SMILES | Group info |
|---|---|---|
| 1 | C1C[C@@H](C([O-])=O)Cc(c2N)c1[nH+]c(c23)cccc3 | 1 |
| 2 | C1C[C@@H](C(=O)N)Cc(c2N)c1[nH+]c(c23)cccc3 | 1 |
| 3 | NCC(=O)[C@H](CC1)Cc(c2N)c1[nH+]c(c23)cccc3 | 1 |
| 4 | COC(=O)[C@H](CC1)Cc(c2N)c1[nH+]c(c23)cccc3 | 1 |
| 5 | CCOC(=O)[C@H](CC1)Cc(c2N)c1[nH+]c(c23)cccc3 | 1 |
| 6 | CNNC(=O)[C@H](CC1)Cc(c2N)c1[nH+]c(c23)cccc3 | 1 |
| 7 | NNC(=O)[C@H](CC1)Cc(c2N)c1[nH+]c(c23)cccc3 | 1 |
| 8 | C1C[C@@H](C(N)=[NH2+])Cc(c2N)c1[nH+]c(c23)cccc3 | 1 |
| 9 | [O-]C(=O)C[C@H](CC1)Cc(c2N)c1[nH+]c(c23)cccc3 | 1 |
| 10 | NC(=O)C[C@H](CC1)Cc(c2N)c1[nH+]c(c23)cccc3 | 1 |
| 11 | NC(=[NH2+])C[C@H](CC1)Cc(c2N)c1[nH+]c(c23)cccc3 | 1 |
| 12 | COC(=N/[H])\C[C@H](CC1)Cc(c2N)c1[nH+]c(c23)cccc3 | 1 |
| 13 | COC(=O)C[C@H](CC1)Cc(c2N)c1[nH+]c(c23)cccc3 | 1 |
| 14 | COC(OC)C[C@H](CC1)Cc(c2N)c1[nH+]c(c23)cccc3 | 1 |
| 15 | CC(C)C[C@H](CC1)Cc(c2N)c1[nH+]c(c23)cccc3 | 1 |
| 16 | CN(C)N[C@H](CC1)Cc(c2N)c1[nH+]c(c23)cccc3 | 1 |
| 17 | N#CN[C@H](CC1)Cc(c2N)c1[nH+]c(c23)cccc3 | 1 (17) |
| 18 | CCOC(=O)[C@H](CC1)Cc(c2N)c1[nH+]c(c23)cccc3 | 1 (16) |
| 19 | NNC(=O)[C@H](CC1)Cc(c2N)c1[nH+]c(c23)cccc3 | 1 |
| 20 | Cn1nccc1NC(=O)[C@H](CC2)Cc(c3N)c2[nH+]c(c34)cccc4 | 1 |
| 21 | c1ccccc1CNC(=O)[C@H](CC2)Cc(c3N)c2[nH+]c(c34)cccc4 | 1 |
| 22 | c1ccccc1CCNC(=O)[C@H](CC2)Cc(c3N)c2[nH+]c(c34)cccc4 | 1 |
| 23 | Cn(c1)ncc1-c(cc2)cc(c23)[nH+]c4c(c3N)CCCC4 | 2 (201) |
| 24 | C1CCCc(c2N)c1[nH+]c(c23)cc(cc3)-c4cncnc4 | 2 (203) |

| | | |
|---|---|---|
| 25 | C1CCCc(c2N)c1[nH+]c(c23)cc(cc3)-c4ccccc4 | 2 |
| 26 | C1CCCc(c2N)c1[nH+]c(c23)cc(cc3)-c4cccnc4 | 2 (214) |
| 27 | c1occc1-c(cc2)cc(c23)[nH+]c4c(c3N)CCCC4 | 2 (212) |
| 28 | n1[nH]ccc1-c(cc2)cc(c23)[nH+]c4c(c3N)CCCC4 | 2 (204) |
| 29 | N#Cc1ccc(cc1)-c(cc2)cc(c23)[nH+]c4c(c3N)CCCC4 | 2 (205) |
| 30 | c1nc(C)ncc1-c(cc2)cc(c23)[nH+]c4c(c3N)CCCC4 | 2 |
| 31 | FC(F)(F)Oc(cc1)ccc1-c(cc2)cc(c23)[nH+]c4c(c3N)CCCC4 | 2 (206) |
| 32 | C1CCCc(c2N)c1[nH+]c(c23)cc(cc3)-c4ccc(O)cc4 | 2 |
| 33 | C1CCCc(c2N)c1[nH+]c(c23)cc(cc3)-c4ccc(F)cc4 | 2 (209) |
| 34 | C1CCCc(c2N)c1[nH+]c(c23)cc(cc3)-c4ccc(N)cc4 | 2 |
| 35 | NC(=O)c1ccc(cc1)-c(cc2)cc(c23)[nH+]c4c(c3N)CCCC4 | 2 |
| 36 | [O-]C(=O)c1ccc(cc1)-c(cc2)cc(c23)[nH+]c4c(c3N)CCCC4 | 2 |
| 37 | COc(cc1)ccc1-c(cc2)cc(c23)[nH+]c4c(c3N)CCCC4 | 2 |
| 38 | C1CCCc(c2N)c1[nH+]c(c23)cc(cc3)-c4c(F)cccc4 | 2 (208) |
| 39 | CSc(cc1)ccc1-c(cc2)cc(c23)[nH+]c4c(c3N)CCCC4 | 2 (210) |
| 40 | FC(F)(F)c1ccc(cc1)-c(cc2)cc(c23)[nH+]c4c(c3N)CCCC4 | 2 (211) |
| 41 | c1c[nH]c(c12)cc(cc2)-c(cc3)cc(c34)[nH+]c5c(c4N)CCCC5 | 2 |
| 42 | C1CCCc(c2N)c1[nH+]c(c23)cc(cc3)-c4ccc(Cl)cc4 | 2 |
| 43 | C1CCCc(c2N)c1[nH+]c(c23)cc(cc3)-c4ccc(Br)cc4 | 2 |
| 44 | C[C@@H]1C[C@@H](C1)c(cc2)cc(c23)[nH+]c4c(c3N)CCCC4 | 2 |
| 45 | CCOC(=O)[C@H](CC1)Cc(c2N)c1[nH+]c(c23)cc(Br)cc3 | 3 (5) |
| 46 | CCOC(=O)[C@H](CC1)Cc(c2N)c1[nH+]c(c23)cc(cc3)-c4ccn(n4)C | 3 (10) |
| 47 | CNC(=O)[C@H](CC1)Cc(c2N)c1[nH+]c(c23)cc(cc3)-c4ccn(n4)C | 3 (14) |
| 48 | C1C[C@@H](C([O-])=O)Cc(c2N)c1[nH+]c(c23)cc(cc3)-c4ccn(n4)C | 3 |
| 49 | Cn(n1)ccc1-c(cc2)cc(c23)[nH+]c4c(c3N)C[C@@H](CC4)N[NH3+] | 3 |
| 50 | Cn(n1)ccc1-c(cc2)cc(c23)[nH+]c4c(c3N)C[C@@H](CC4)OC | 3 |
| 51 | CO[C@H](CC1)Cc(c2N)c1[nH+]c(c23)cc(cc3)-c4cc[nH]n4 | 3 |

| | | |
|---|---|---|
| 52 | C1C[C@@H](C([O-])=O)Cc(c2N)c1[nH+]c(c23)cc(cc3)-c4cc[nH]n4 | 3 |
| 53 | [O-]C(=O)C[C@H](CC1)Cc(c2N)c1[nH+]c(c23)cc(cc3)-c4cc[nH]n4 | 3 |
| 54 | C1C[C@@H](C)Cc(c2N)c1[nH+]c(c23)cc(cc3)-c4cc[nH]n4 | 3 |
| 55 | CCOC(=O)[C@H](CC1)Cc(c2N)c1[nH+]c(c23)cc(Cl)cc3 | 3 |
| 56 | CCOC(=O)[C@H](CC1)Cc(c2N)c1[nH]c(c23)cc(=O)cc3 | 3 |
| 57 | CCOC(=O)[C@H](CC1)Cc(c2N)c1[nH+]c(c23)cc(cc3)OC | 3 |
| 58 | CCOC(=O)[C@H](CC1)Cc(c2N)c1[nH+]c(c23)cc(cc3)-c4ccc(F)cc4 | 3 (107) |
| 59 | CCOC(=O)[C@H](CC1)Cc(c2N)c1[nH+]c(c23)cc(cc3)-c4ccccc4 | 3 |
| 60 | CCOC(=O)[C@H](CC1)Cc(c2N)c1[nH+]c(c23)cc(cc3)-c4cnc[nH]4 | 3 |
| 61 | NNC(=O)[C@H](CC1)Cc(c2N)c1[nH+]c(c23)cc(Br)cc3 | 3 (8) |
| 62 | NNC(=O)[C@H](CC1)Cc(c2N)c1[nH+]c(c23)cc(Cl)cc3 | 3 |
| 63 | NNC(=O)[C@H](CC1)Cc(c2N)c1[nH+]c(c23)cc(cc3)-c4ccccc4 | 3 |
| 64 | NNC(=O)[C@H](CC1)Cc(c2N)c1[nH+]c(c23)cc(cc3)-c4cnc[nH]4 | 3 |
| 65 | NNC(=O)[C@H](CC1)Cc(c2N)c1[nH+]c(c23)cc(cc3)-c4cncnc4 | 3 |
| 66 | CNC(=O)[C@H](CC1)Cc(c2N)c1[nH+]c(c23)cc(cc3)-c4cncnc4 | 3 |
| 67 | CNC(=O)[C@H](CC1)Cc(c2N)c1[nH+]c(c23)cc(cc3)-c4cnc[nH]4 | 3 |
| 68 | CNC(=O)[C@H](CC1)Cc(c2N)c1[nH+]c(c23)cc(Br)cc3 | 3 (13) |
| 69 | CNC(=O)[C@H](CC1)Cc(c2N)c1[nH+]c(c23)cc(Cl)cc3 | 3 |
| 70 | CNC(=O)[C@H](CC1)Cc(c2N)c1[nH+]c(c23)cc(cc3)-c4cncnc4 | 3 |
| 71 | CNC(=O)[C@H](CC1)Cc(c2N)c1[nH+]c(c23)cc(cc3)-c4ncc[nH]4 | 3 |
| 72 | C1C[C@@H](C(=O)N)Cc(c2N)c1[nH+]c(c23)cc(Br)cc3 | 3 |
| 73 | C1C[C@@H](C(=O)N)Cc(c2N)c1[nH+]c(c23)cc(Cl)cc3 | 3 |
| 74 | C1C[C@@H](C(=O)N)Cc(c2N)c1[nH+]c(c23)cc(cc3)-c4ccccc4 | 3 |
| 75 | C1C[C@@H](C(=O)N)Cc(c2N)c1[nH+]c(c23)cc(cc3)-c4cnc[nH]4 | 3 |

**Table S1:** 75 designed tacrine derived MTDLs, their SMILEs notation and the information about the group to which they belong to. The numbers shown in parentheses indicate the numbers used in our patent applications (WO 2019/207604 A1, IPO-201841015699).

| Name | MW (≤450Da) | HBD (≤37 | HBA (≤3) | LogP (≤5) | CNS (≥0) | LogBB (-1.2-1.2) | MDCK (>500nm/s) | Rotor (<8) | PSA (<90 Å) | %HOA (61-100) |
|---|---|---|---|---|---|---|---|---|---|---|
| 5 | 349 | 1.5 | 4 | 3.47 | 0 | -0.43 | 1430 | 3 | 69 | 100 |
| 8 | 335 | 4.5 | 5 | 1.41 | -2 | -1.11 | 209 | 3 | 102 | 76 |
| 10 | 350 | 1.5 | 5.5 | 3.71 | -1 | -0.86 | 387 | 3 | 85 | 100 |
| 13 | 334 | 2.5 | 4.5 | 2.06 | -1 | -0.36 | 1312 | 2 | 72 | 89 |
| 14 | 335 | 2.5 | 6 | 2.31 | -1 | -0.78 | 354 | 2 | 87 | 88 |
| 16 | 270 | 1.5 | 4 | 2.91 | 0 | -0.59 | 539 | 3 | 68 | 100 |
| 17 | 256 | 4.5 | 5 | 0.88 | -2 | -1.25 | 79 | 3 | 102 | 73 |
| 107 | 364 | 1.5 | 4 | 4.76 | 0 | -0.65 | 980 | 4 | 69 | 100 |
| 201 | 278 | 1.5 | 3.5 | 3.27 | 0 | -0.29 | 801 | 1 | 52 | 100 |
| 203 | 276 | 1.5 | 5 | 2.38 | 0 | -0.59 | 412 | 2 | 60 | 93 |
| 204 | 264 | 2.5 | 3.5 | 2.49 | -1 | -0.53 | 403 | 1 | 63 | 94 |
| 205 | 299 | 1.5 | 3.5 | 3.48 | -1 | -0.87 | 286 | 3 | 60 | 100 |
| 206 | 358 | 1.5 | 2 | 5.34 | 1 | 0.13 | 7326 | 3 | 42 | 100 |
| 208 | 292 | 1.5 | 2 | 4.44 | 1 | 0.002 | 2489 | 2 | 34 | 100 |
| 209 | 292 | 1.5 | 2 | 4.43 | 1 | 0.03 | 2840 | 2 | 34 | 100 |
| 210 | 320 | 1.5 | 2.5 | 4.84 | 0 | -0.07 | 2677 | 3 | 34 | 100 |
| 211 | 342 | 1.5 | 2 | 5.19 | 1 | 0.18 | 6936 | 2 | 34 | 100 |
| 212 | 264 | 1.5 | 2.5 | 3.41 | 1 | 0.02 | 1570 | 1 | 43 | 100 |
| 214 | 275 | 1.5 | 3.5 | 3.30 | 0 | -0.34 | 811 | 2 | 47 | 100 |
| tacrine | 198 | 1.5 | 2 | 2.57 | 1 | 0.04 | 1570 | 1 | 34 | 100 |

**Table S2:** Important physico-chemical properties predicted using QikProp for the selected tacrine derived MTDLs. Abbreviations: MW: Molecular weight; HBD: Hydrogen bond donors; HBA: Hydrogen bond acceptors; MDCK: Mandin-Darby canine kidney; PSA: Polar surface area; %HOA: % of human oral absorption. The recommended range for CNS active drugs is given in parenthesis.

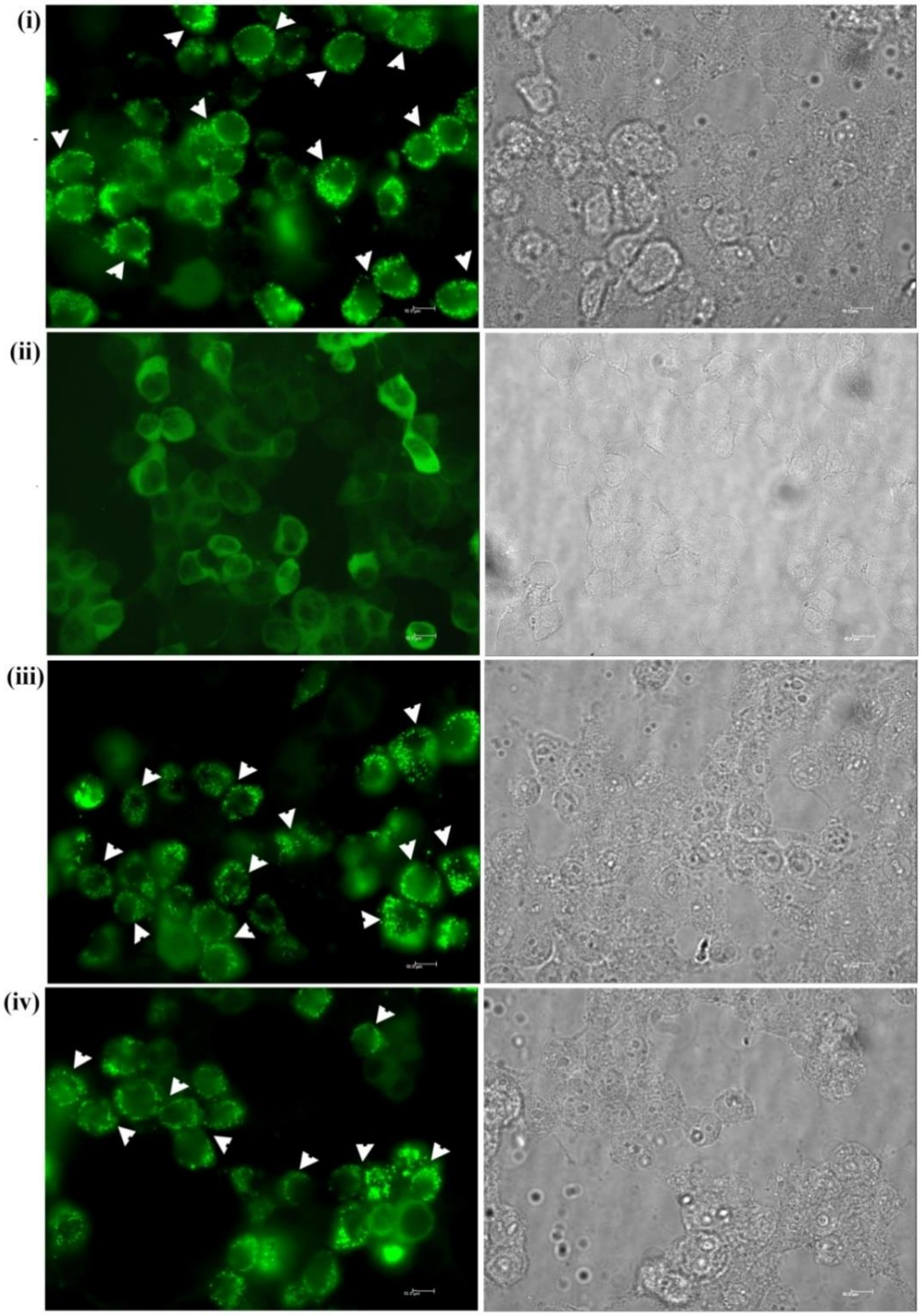


**Figure S1:** Tacrine derived MTDLs do not affect α-CaMKII-GluN2B interaction. Assays were performed on HEK-293 cells that stably express GFP-α-CaMKII and MLS-NR2B (a portion of GluN2B having α-CaMKII binding motif with MLS tag) without functional NMDAR. α-CaMKII activated by $Ca^{2+}$ influx through the ionophore, ionomycin, binds to MLS-NR2B and gives rise to punctae (i) which is not seen in the absence of $Ca^{2+}$ (ii). The treatment of MTDLs such as 208 (iii) and 211 (iv) along with ionomycin and $Ca^{2+}$ did not cause any significant reduction in the extent of punctae formation.

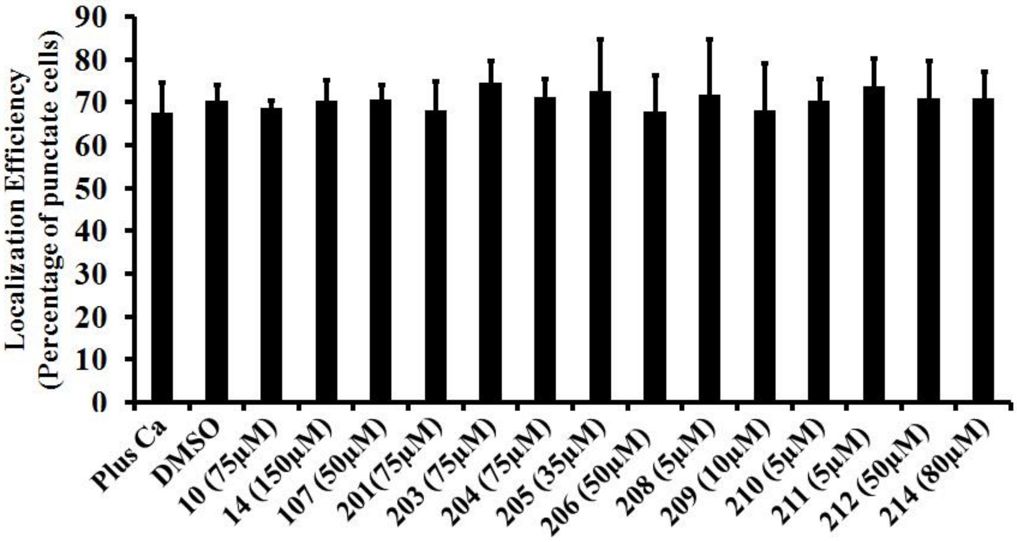


**Figure S2:** Quantitation of punctae formation in presence of selected tacrine derived MTDLs along with ionomycin and $Ca^{2+}$. HEK-293 cells stably expressing GFP-α-CaMKII and MLS-NR2B without functional NMDAR were subjected to treatment with ionomycin and $Ca^{2+}$ in presence of the indicated MTDL as described in Figure S1. Subsequently the punctate cell count was quantitated as mentioned in Methods section 'NMDAR activity assay'.

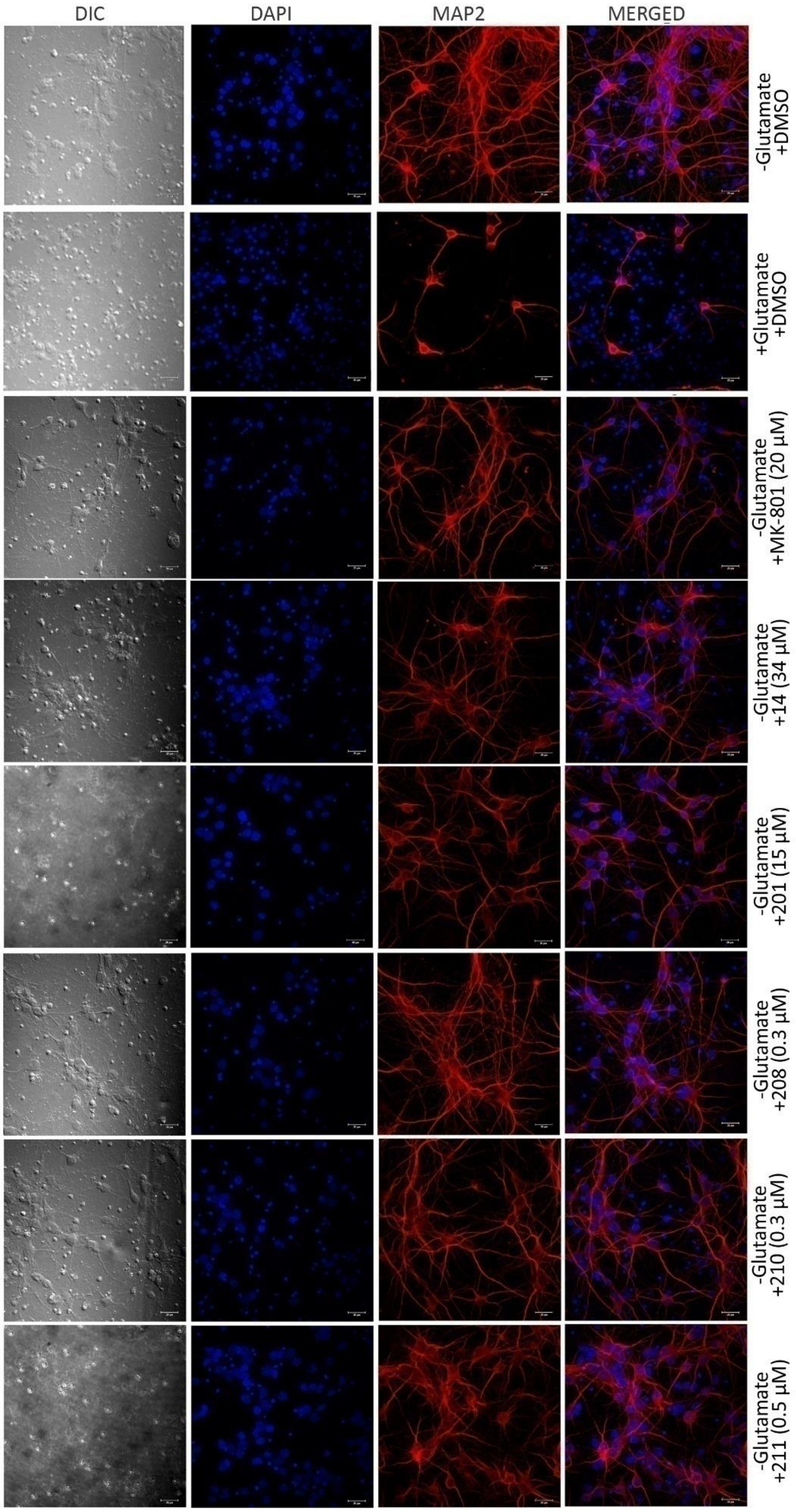
DIC
DAPI
MAP2
MERGED
-Glutamate +DMSO
+Glutamate +DMSO
-Glutamate +MK-801 (20 μM)
-Glutamate +14 (34 μM)
-Glutamate +201 (15 μM)
-Glutamate +208 (0.3 μM)
-Glutamate +210 (0.3 μM)
-Glutamate +211 (0.5 μM)

**Figure S3:** Representative images of primary cortical neurons immunostained for the neuronal protein MAP2 (red) with nuclei counterstained using DAPI (blue) after treatment in the absence or presence of 100 µM glutamate for 1 hour. The cells subjected to treatment in presence of selected MTDLs (14, 201, 208, 210 and 211) in the absence of glutamate are also shown. Since DMSO was used to dissolve the MTDLs, DMSO was included in the treatments without MTDLs also.

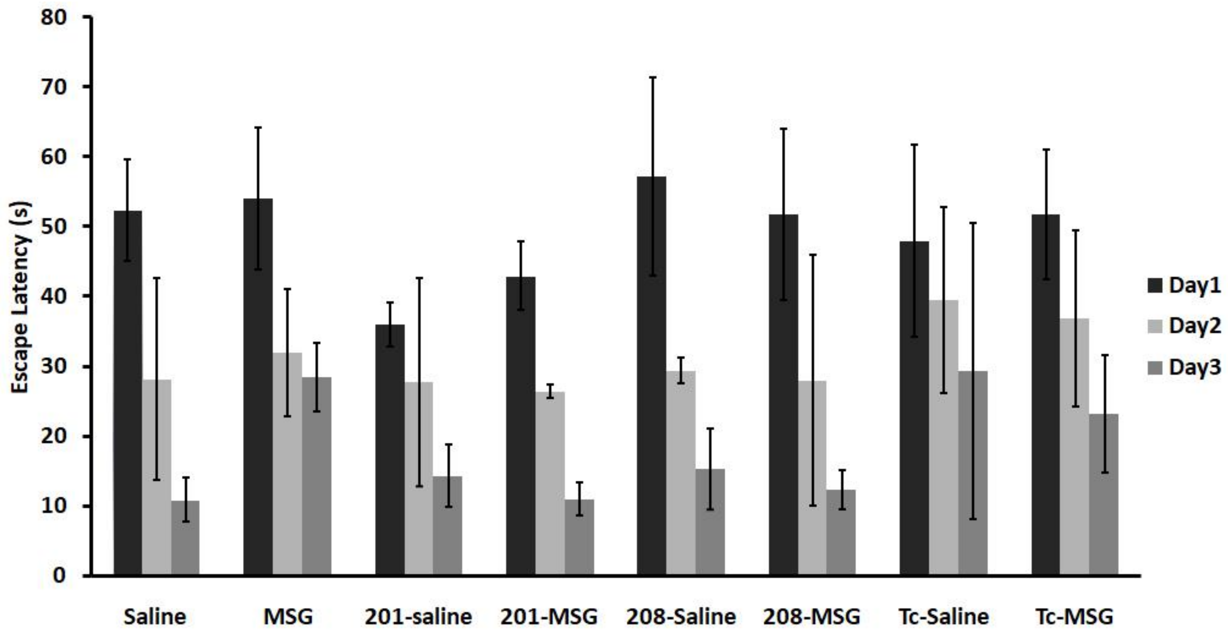


**Figure S4:** Effect of tacrine derived MTDLs (201 and 208) on MSG induced behavioral impairment determined by MWM test. Escape latencies to reach the platform from Day1 to Day3 are given. Five trials for each day were averaged for each animal. Data are presented as the mean ± SD, n=3. Data for Day3 is the same as that presented in Figure 4A.
